# Seasonality and the persistence of vector-borne pathogens

**DOI:** 10.1101/2022.12.13.520207

**Authors:** Van Hai Khong, Philippe Carmona, Sylvain Gandon

## Abstract

Many vector-borne diseases are affected by the seasonality of the environment. Yet, seasonality can act on distinct steps of the life-cycle of the pathogen and it is often difficult to predict the influence of the periodic fluctuations of the environment on the persistence of vector-borne pathogens. Here we analyse a general vector-borne disease model and we account for periodic fluctuations of different components of the pathogen’s life-cycle. We develop a perturbation analysis framework to obtain useful approximations to evaluate the overall consequences of seasonality on the persistence of pathogens. This analysis reveals when seasonality is expected to increase or to decrease pathogen persistence. We show that seasonality in vector density or in the biting rate of the vector can have opposite effects on persistence and we provide a useful biological explanation for this result based on the covariance between key compartments of the epidemiological model. This framework could be readily extended to explore the influence of seasonality on other components of the life cycle of vector-borne pathogens.

**Significance statement:** Does seasonality increase or decrease the persistence of vector-borne diseases? The devil is in the details and our analysis shows that the effect of seasonality depends on which pathogen traits are affected by seasonality. We highlight the contrasting effects of seasonality in the abundance and/or in the biting rate of the vector on pathogen persistence.

## 1 Introduction

The ability of a pathogen to spread in a fully susceptible host population is governed by its basic reproduction ratio *R*_0_ which measures the number of secondary cases produced by a typical infected case. The pathogen will spread and induce an epidemic if and only if *R*_0_ > 1 (and it will rapidly go extinct if *R*_0_ < 1). Hence the basic reproduction ratio provides a way to evaluate the epidemic potential of different pathogens and constitutes a key epidemiological quantity to control infectious diseases. The basic reproduction ratio of the simplest epidemiological model with a single compartment of infected hosts takes a very simple and intuitive form: *R*_0_ = *β/γ* (i.e., the transmission rate *β* times the duration of infection 1/*γ*). The next generation matrix (a per-generation way to compute population dynamics) can be used to compute *R*_0_ when multiple compartments (e.g., different infectious stages, different hosts) are required to describe the life cycle of the pathogen in a constant environment where transitions rates are fixed [5]. Indeed, in a constant environment, a per-generation way of measuring population growth is appropriate because the timing of the new infection does not matter.

But things become more complicated in fluctuating environments where the timing of new infections matter. For instance, many studies have shown that a periodically changing environments may affect the *R*_0_ [3, 8, 9, 20]. The analysis of these models is more challenging and does not always yield consistent results regarding the qualitative effects of seasonality on *R*_0_. For instance, periodic fluctuations in the density of mosquito vectors can reduce *R*_0_ [3] while a fluctuation in the transmission rate yields higher *R*_0_ in another vector-borne model [20]. Why these different models yield opposite conclusions? Can we build up an intuitive and biological understanding of the behaviour of these complex time-varying models? We try to answer these questions in the following with the analysis of simple periodic models of vector-borne pathogens. We derive different threshold quantities for the persistence of the pathogen that allow us to discuss the effects of periodic fluctuations of various components of the pathogen’s life cycle.

## 2 The model

Let us consider the spread of a vector-borne pathogen which requires an explicit description of the dynamics of the infection among human hosts and mosquito vectors. We use a classical model of vector-borne transmission [2] that tracks the densities of four types of hosts (uninfected and infected humans, uninfected and infected vectors) which yields the following system of ordinary differential equations (the parameters of the model are described in Table 1):

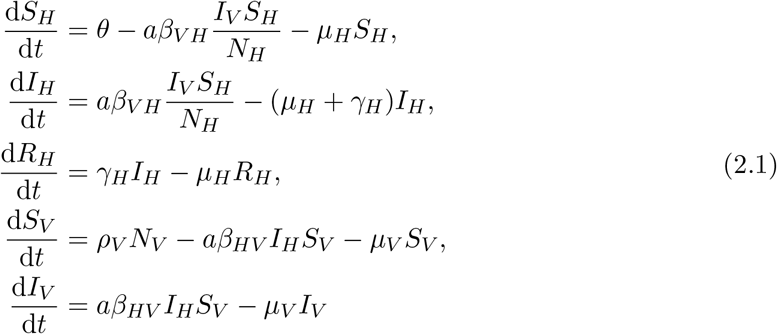

For the sake of simplicity we assume that the total density of the human population *S_H_* + *I_H_* + *R_H_* = *N_H_* remains constant which implies that *θ* = *μ_H_*(*S_H_* + *I_H_* + *R_H_*).

**Table 1:** Parameters and variables of the models

| Symbol | Description |
| --- | --- |
| Parameters |  |
| $a$ | biting rate of vectors |
| $\beta_{VH}$ | rate of transmission from an infected vector to a susceptible host |
| $\beta_{HV}$ | rate of transmission from an infected host to a susceptible vector |
| $\rho_V$ | growth rate of the vector |
| $\mu_V$ | death rate of the vector |
| $\mu_H$ | death rate of the host |
| $\gamma_H$ | recovery rate of the infected host |
| Variables |  |
| $\theta$ | the periodic influx of susceptible hosts |
| $S_H$ | density of susceptible hosts |
| $I_H$ | density of infected hosts |
| $R_H$ | density of recovered hosts |
| $N_H$ | density of the total host population: $N_H = R_H + I_H + S_H$ |
| $S_V$ | density of susceptible vectors |
| $I_V$ | density of infected vectors |
| $N_V$ | total vector population density: $N_V = S_V + I_V$ |

In a constant environment we assume that the parameters that govern the life cycle of the vector do not vary with time and if the fecundity of the vector compensates exactly its mortality (i.e., *ρ_V_* = *μ_V_*) the total density of the vector population *S_V_* + *I_V_* = *N_V_* is also constant. In this scenario the basic reproduction ratio is readily derived from the linearisation of the above system at the disease free equilibrium (*I_V_* = *I_H_* = *R_H_* = 0, *S_H_* = *N_H_, S_V_* = *N_V_*) which yields ([8]):

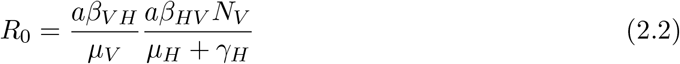

In other words, the *R*_0_ can be interpreted as the expected number of human hosts infected by an infected vector during its lifetime (the first ratio in (2.2)) times the expected number of vectors infected by an infected host throughout the duration of its infection (the second ratio in (2.2)).

In a fluctuating environment several parameters of the pathogen may vary with time [15, 19] and these variations are likely to affect *R*_0_. Several previous studies have developed ways to compute *R*_0_ in periodic environments [3, 8, 9, 20]. Seasonality is usually modelled through the use the T-periodic function *f*(*t*) = cos(2*πt*/*T*) which may affect different biological processes and thus different parameters in the dynamical system (2.1). Bacaer analysed a model of malaria transmission when the density of the mosquito population fluctuates periodically with *N_V_*(*t*) = *N_V_*(1 + *ϵf*(*t*)) [3]. This analysis combines a Fourier decomposition of *N_V_*(*t*) with a perturbation analysis for small *ϵ* to compute an approximation of *R*_0_ as a function of *ϵ*. This approach yields a particularly useful approximation for the effect of seasonality:

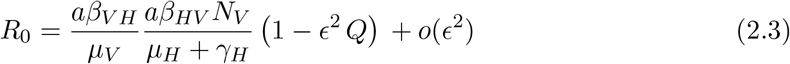

where 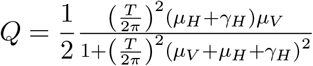 This expression shows that, since *Q* > 0, the periodic fluctuations of the densities of mosquitoes tend to decrease *R*_0_ and thus to limit the persistence of malaria. Yet, it is difficult to understand *why* this is the case.

An alternative approach to handle the analysis of periodic system is to use Floquet theory which is based on the analysis of the linearization of (2.1) which yields:

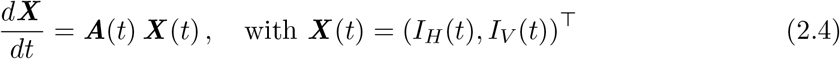

with 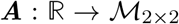 a continuous matrix valued *T* periodic function and 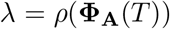 the spectral radius of the monodromy matrix of system (2.4), that is

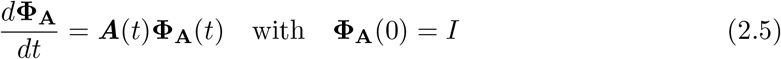

In other words, λ is the asymptotic growth rate of the epidemic in the initial phase of the epidemic and if λ > 1 the pathogen density grows exponentially and 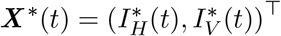 is the (unique up to a constant) positive vector solution of (2.4) such that 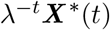 has period *T*. It is obtained by letting ***X***\*(0) be the positive eigenvector associates to the monodromy matrix and its spectral radius: 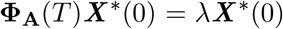. The vector λ^-*t*^***X***\* may be thought as the stable composition of the pathogen population in the periodic environment, for the linearized model (2.5).

Crucially, as pointed out by Heesterbeek and Roberts [8, 9], this formalism yields another interesting quantity which is akin to the original definition of *R*_0_ as it provides the expected number of secondary cases :

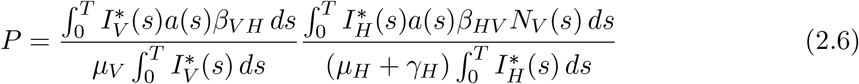

As in (2.2) the first ratio of (2.6) refers to the expected number of human hosts infected by an infected vector during its lifetime, while the second ratio of (2.6) refers to the expected number of vectors infected by an infected human throughout the duration of its infection. The integral over one period of the fluctuation accounts for the fluctuations of the different parameters of the pathogen’s life-cycle. This quantity can also be expressed in terms of the covariances between different key epidemiological variables:

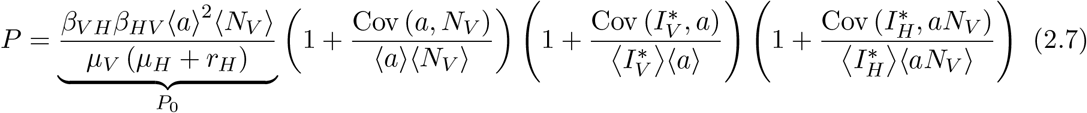

As expected, in the absence of temporal fluctuations all the covariances vanish and *P*_0_ is equal to *R*_0_ given in (2.2). We will see in the following that this expression of *P* can be particularly useful to understand the effects of seasonality.

Thanks to the works of Heesterbeek and Roberts [8, 9] (see also Wang and Zhao [20]) we thus have three threshold parameters *R*_0_, λ and *P* that can be used to determine the ability of the pathogen to invade the population. Indeed, even if these three quantities are not equivalent, they are all equal when the pathogen reaches the threshold value *R*_0_ = 1. Since the signs of *R*_0_ – 1, λ – 1 and *P* – 1 are the same we can analyse the effect of seasonality on persistence of the pathogen population using these three quantities around *R*_0_ = λ = *P* = 1. Note, however, that the effect of seasonality we discuss in this limit remain qualitatively robust for other values of *R*_0_.

In the following we study the effect of seasonality on the stability of the disease-free equilibrium through the effect of *ϵ* which captures the magnitude of the influence of seasonality of distinct parameters of the model. We use a perturbation analysis of λ for small values of *ϵ* at λ = 1. First, we contrast the effect of seasonality on various quantities that affect the vector population and the transmission of the disease. We show that the effect of seasonality on λ depends on which trait is affected by the fluctuations of the environment. Second, we use the quantity *P* to provide a biological interpretation of these effects of seasonality in terms of a covariance between different dynamical variables.

## 3 Results

Seasonality is known to affect various components of the life-cycle of vector-borne pathogens [15]. In the following we assume that seasonal variations may act directly on the density of the vector population and/or on the biting rate of mosquitoes. More specifically we use the following T-periodic functions with *f*(*t*) = cos(2*πt/T*):

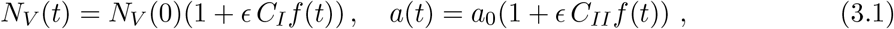

where *C_I_* ≥ 0 and *C_II_* ≥ 0 control the magnitude of the effect of seasonality on *N_V_*(*t*) and *a*(*t*), respectively. This model allows to account for the effects of seasonality on multiple traits of the pathogen. In the following, we contrast two extreme scenarios: (i) model *I* when *C_I_* = 1 and *C_II_* = 0 and (ii) model *II* when *C_I_* = 0 and *C_II_* = 1.

We derive an approximation for λ for small values of *ϵ* (see Supplementary Information):

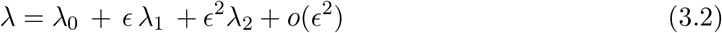

with λ_0_ = 1 because we focus on the persistence of the pathogen population and where λ_1_ and λ_2_ refer to the first and second order effects of seasonality with:

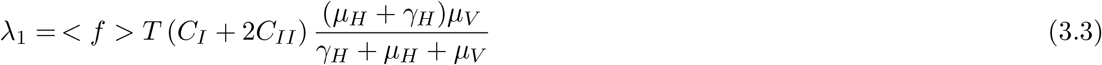

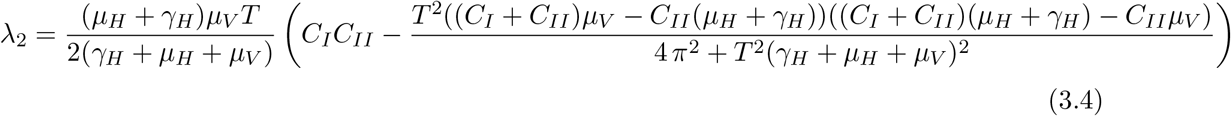

### 3.1 First order effect

Seasonality has a first order effect on the stability of the disease-free equilibrium as soon as < *f* >≠ 0. For instance, consider a simple square function where *f*(*t*) = 1 for *t* ∈ [0, (1 – *W*)*T*] and *f*(*t*) = –1 for *t* ∈ [(1 – *W*)*T, T*] (with 0 < *W* < 1) as illustrated in **Figure 1**. The parameter *W* governs the duration of the “winter”. Higher values of *W* yield longer winter seasons where the total density of vectors is reduced (model *I*) or when the biting rate of the vector is reduced (model *II*). **Figure 1** shows that longer winters (i.e., *W* > 0.5) yield < *f* >< 0 and, in this case, more seasonality (i.e., higher values of *ϵ*) reduces pathogen persistence in both models *I* and *II*. In contrast, shorter winters (i.e., *W* < 0.5) yield < *f* >> 0 and, in this case, more seasonality increases pathogen persistence in both models *I* and *II*. Note that the slope of λ at *ϵ* = 0 is λ_1_, which is twice larger for model *II* than for model *I*, in agreement with equation (3.3).

**Figure 1:**
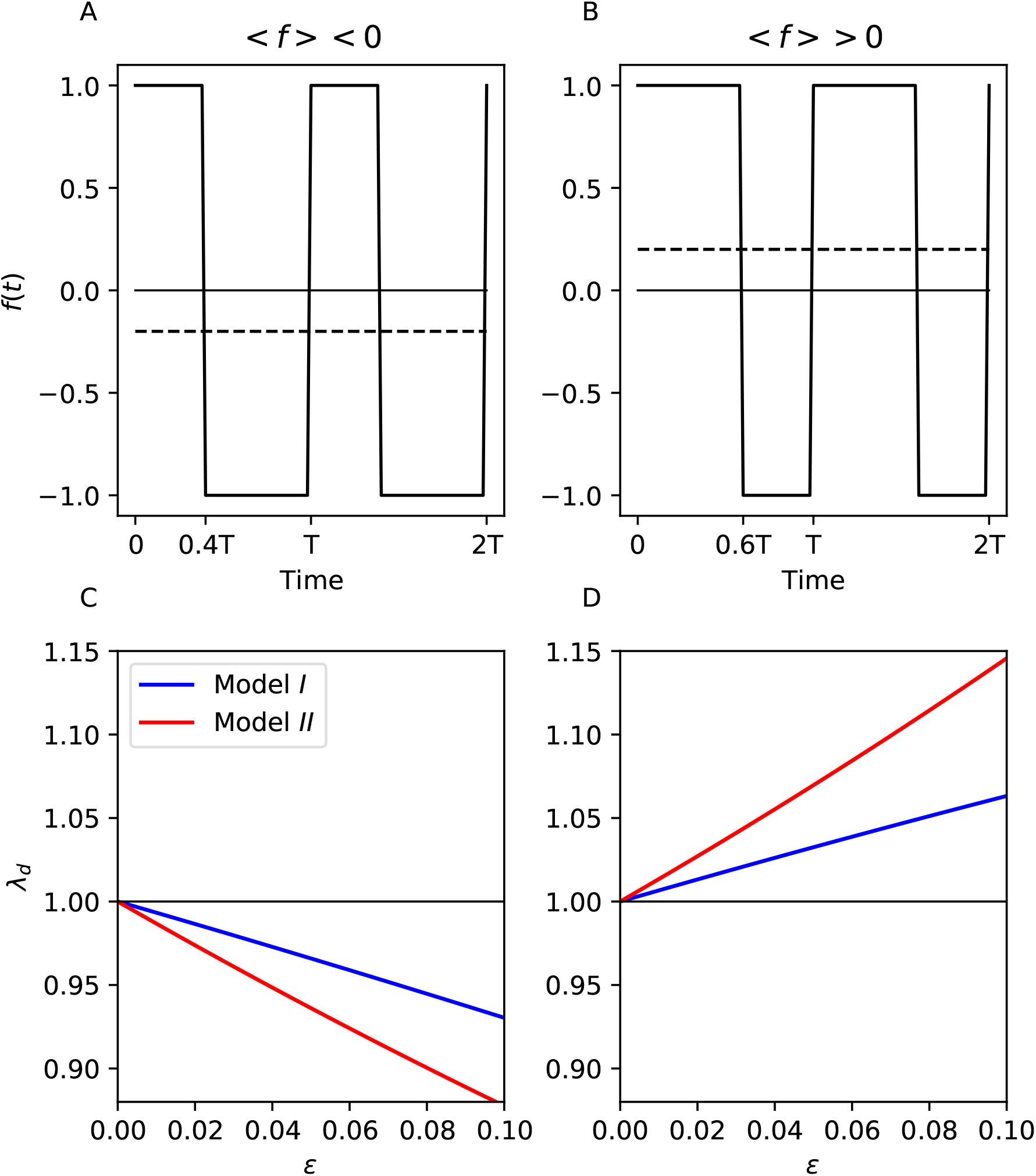
The first order effect of seasonality drives the stability of the disease-free equilibrium when < *f* >≠ 0. In **(A**) We assume *x* = 0.4 and thus 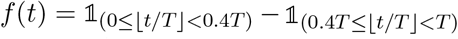. In **(B**) we assume *x* = 0.6 and thus 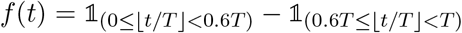. In **(C**) and **(D**) we plot the effect of seasonality (i.e., *ϵ*) on λ_*ϵ*_ for the two scenarios **(A**) and **(B**). In model *I*, *C_I_* = 1, *C_II_* = 0 so that *N_V_* = 1 + *ϵf*(*t*), *a*(*t*) = 1. In model *II*, *C_I_* = 0, *C_II_* = 1, so that *N_V_*(*t*) = 1, *a*(*t*) = 1 + *ϵf*(*t*). Parameters values: *T* = 5, *β_HV_* = 1, *β_VH_* = 2, *μ_H_* = 0.5, *μ_V_* = 1, *γ_H_* = 1.5.

However, this first order effect of seasonality vanishes when < *f* >= 0. In particular, < *f* >= 0 for the classical scenarios we considered where *f* (*t*) = cos(2*πt/T*). In this case we thus need to examine the second order effect of seasonality in (3.4).

### 3.2 Second order effect

Next, we analyse the second order effect of seasonality on λ and, in contrast with the previous section, we show that seasonality has different qualitative effects on λ in models *I* and *II* (Figure (2)).

**Figure 2:**
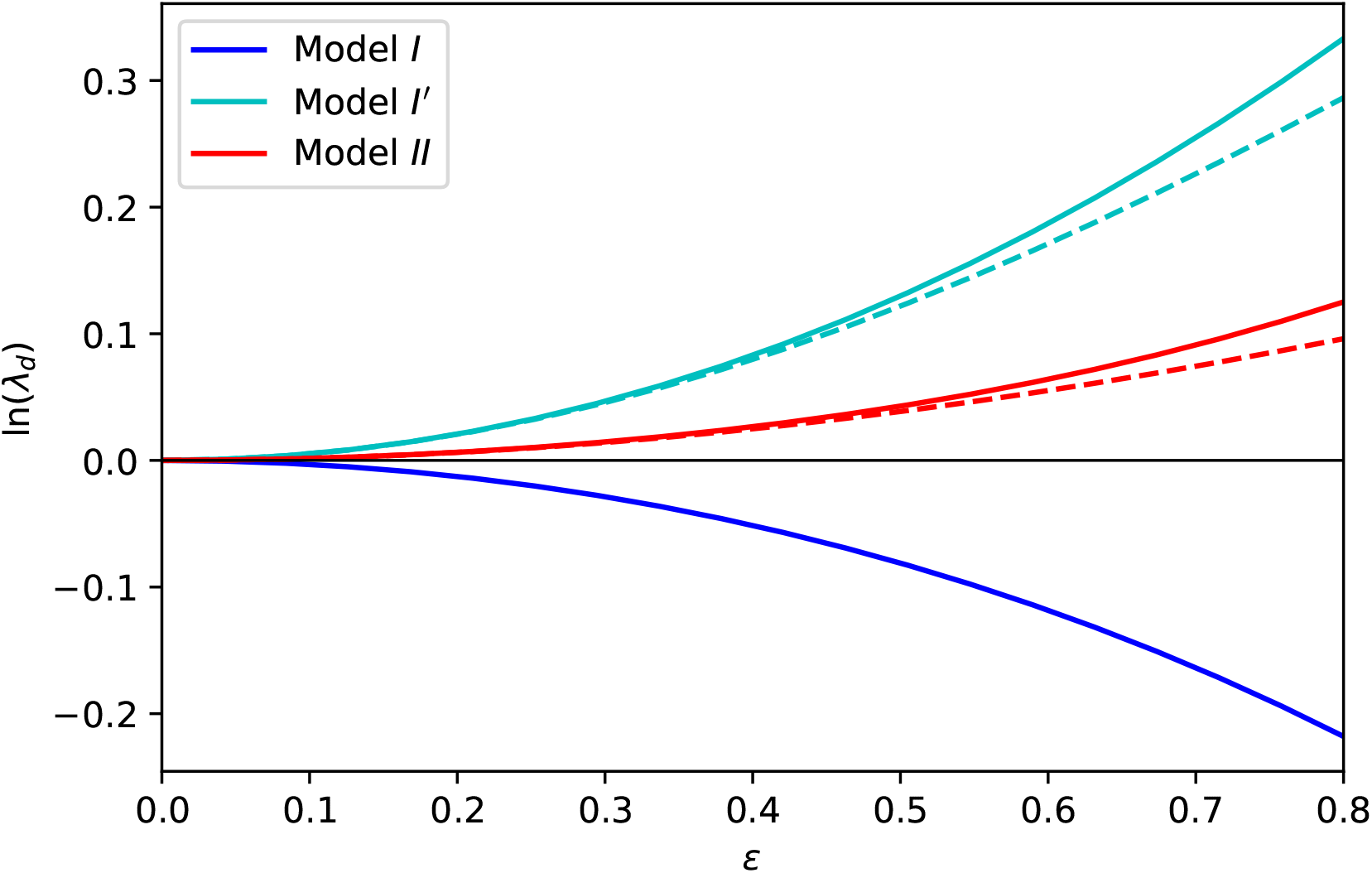
The second order effect of seasonality on λ_*ϵ*_ for three different models. The blue, cyan and red curves represent the value of log λ_*ϵ*_ in models *I, I*′ and *II*, respectively. The solid line indicates exact numerical result while the dashed lines indicates the second order approximation given in equations (3.5), (3.11) and (3.10). In models *I* and *I*′: *C_I_* = 1, *C_II_* = 0. In model *II*: *C_I_* = 0, *C_II_* = 1. Parameter values: *T* = 5, *f*(*t*) = cos(2*πt/T*), *μ_H_* = 0.5, *μ_V_* = 1, *γ_H_* = 1.5, *β_HV_* = 1, *β_VH_* = 2.

#### 3.2.1 Fluctuations in vector density

In model *I* we have *C_I_* = 1 and *C_II_* = 0 and equation (3.2) reduces to

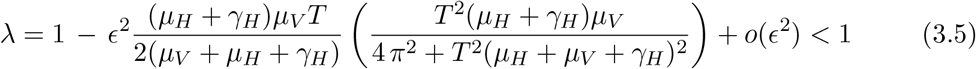

We thus recover the effect of seasonality discussed by Bacaer [3] where fluctuations of vector densities reduce the persistence of the pathogen.

But we can use the threshold quantity *P* to gain some insight in the understanding of the qualitative effect of the fluctuations of vector densities. In particular, we can use (2.6) to show that the stability of the disease-free equilibrium is governed by the covariance between 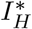 and *N_V_*. In the Supplementary Information we derive an approximation of this covariance which shows that this covariance is always negative, which implies that *P* < 1 and thus that λ < 1 (3.6):

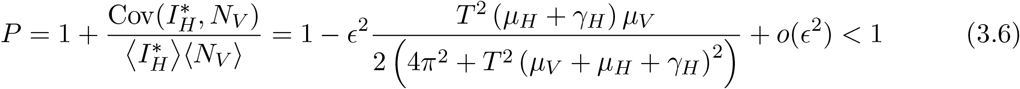

Since the effect of seasonality is governed by the sign of the covariance between 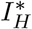 and *N_V_* it is important to understand why this covariance is negative. We expect this covariance to be negative whenever the lag between the fluctuations of *N_V_* and the fluctuations of 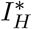 are larger than *T*/4 (the covariance is nil when the lag is exactly equal to *T*/4 which corresponds to a phase lag of π/2). Fluctuations in *N_V_* first drive fluctuation in *I_V_* with a lag approximately equal to *T*/4 (**Figure 4**). Second, these fluctuations in the density of infected vectors drive the fluctuations in the dynamics in *I_H_*. This two-step process yields a lag > *T*/4 between *N_V_* and 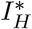, which results in a negative covariance between 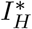 and *N_V_* (see **Figure 4**).

**Figure 3:**
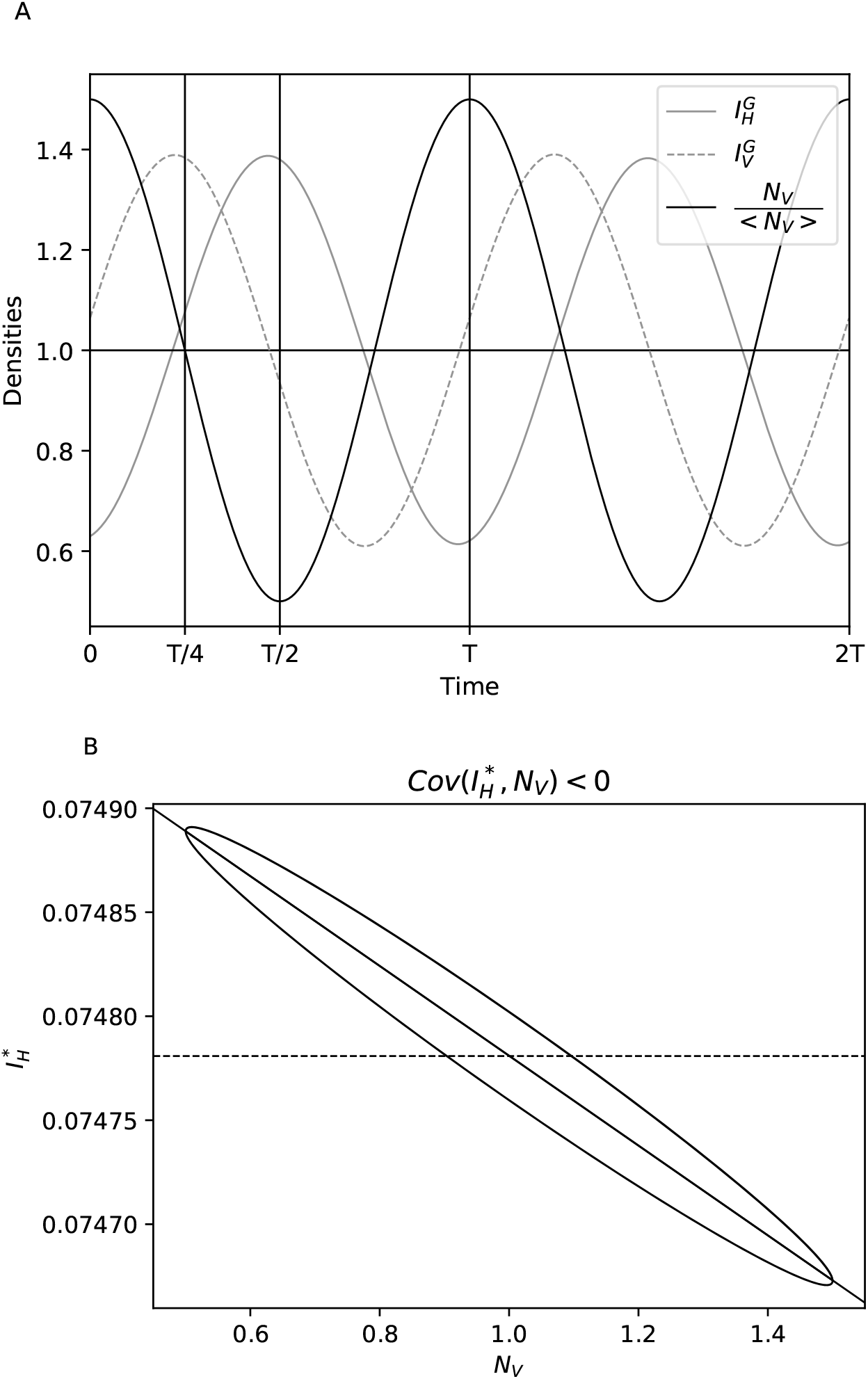
Understanding the covariance between *N_V_* and 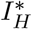 in model *I*. In **(A**), we plot the density of the vector population *N_V_* (full black line), the density of infected vectors *I_V_* (dashed gray line with scale 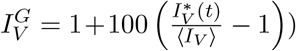, the density of infected hosts *I_H_* (full gray line with scale 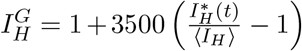. Note that the densities 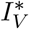 and 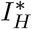 are rescaled using 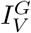 and 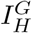 to better visualize the lag with the fluctuations of *N_V_*. The density of the infected vector population 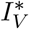 follows the fluctuations of *N_V_* with a lag approximately equal to *T*/4. The density of infected hosts 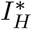 is driven by the fluctuations of infected vectors which yields a larger lag behind the fluctuations of *N_V_* (this lag is close to *T*/2). In **(B**) we present the dynamics of *N_V_* and 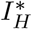 for the same parameter values to show how a lag > *T*/4 results in a negative 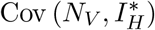. The sign of 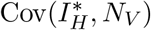 is given by the slope of the regression line (full black line). Parameter values: *a*_0_ = *N_V_*(0) = *β_HV_* = 1, 1 = *β_VH_*, *ϵ* = 0.5, *T* = 1, *μ_V_* = 1, *μ_H_* = 0.01, *γ_H_* = 0.1.

**Figure 4:**
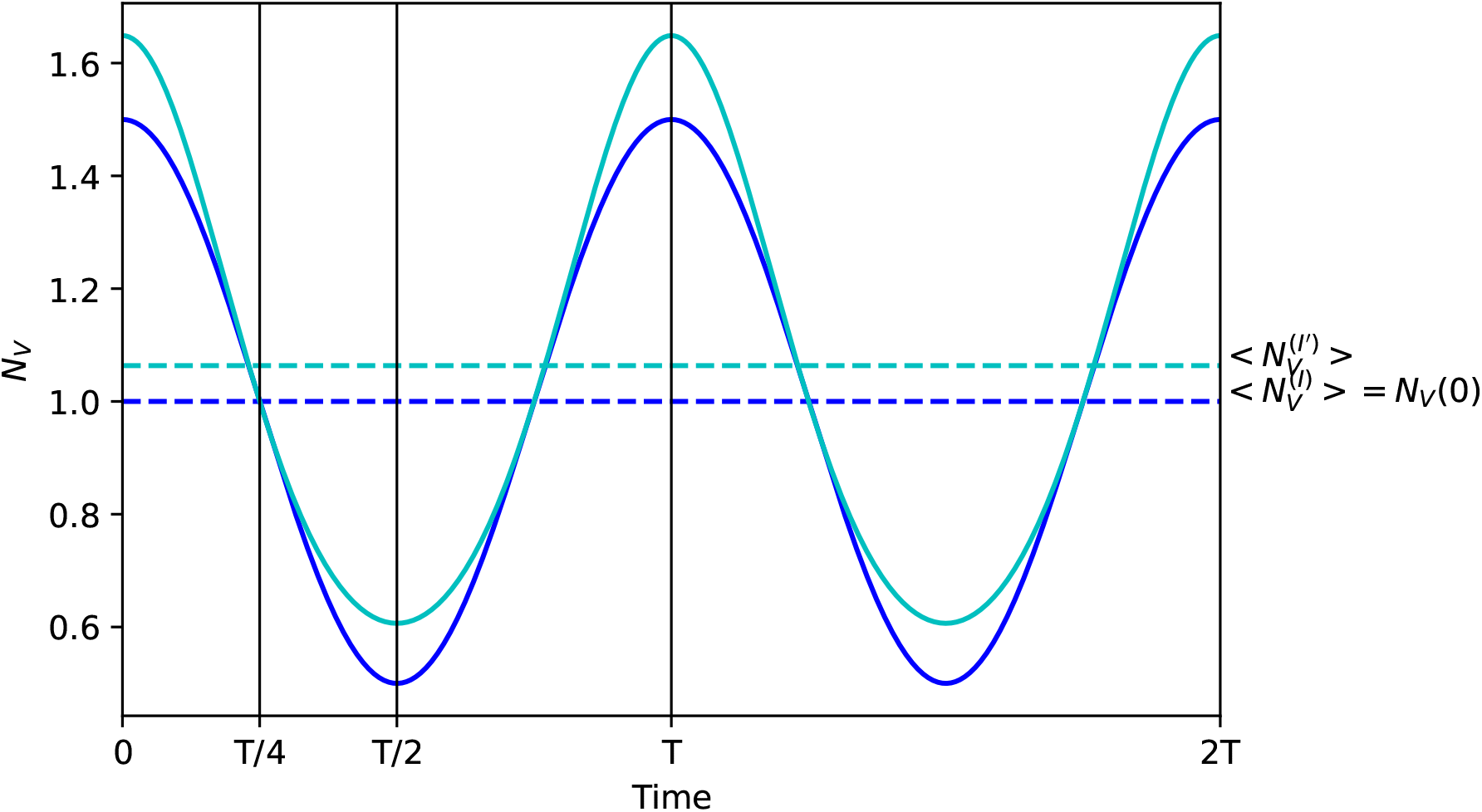
Fluctuations of the density of vectors in models *I* and *I*′. In model *I* (blue curve) we assume *N_V_*(*t*) = 1 + *ϵf*(*t*). In model *I*′ (cyan curve) we assume 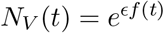. The dashed lines indicate the average value of the density of the vector population and we see that this average value is higher in model *I*′. This effect on the average density of vectors drives the first order effect of seasonality in model *I*′ we illustrate in Figure 2. Parameter values: *ϵ* = 0.5 and *T* = 10, *f*(*t*) = cos(2*πt/T*).

At this stage it is tempting to conclude that any fluctuation in the total density of the vector population always results in the same qualitative effect. But, as we will see in the following, some fluctuation in vector density may have the opposite effect on pathogen persistence. Note that the fluctuations of *N_V_*(*t*) assumed in model *I* (see equation (3.1)) results from specific assumptions regarding the per-capita growth rate of the vector population: 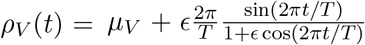. In the following we study another model (model *I*′) where we use a different periodic function to model the fluctuation of the per-capita growth rate of the vector population *ρ_V_*(*t*) = *μ_V_*(1 + *ϵ* cos(2*πt/T*)) as in [9, 13]. This yields the following vector dynamics (see **Figure 4** and Supplementary Information section 4):

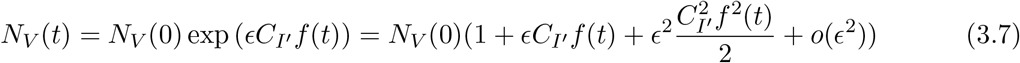

Again the examination of *P* helps understand what is going on. The average value of *N_V_* is affected by seasonality and fluctuations increase the mean density of vectors which feed backs on transmission. Indeed, unlike model *I* where < *N_V_* >= *N_V_*(0), in model 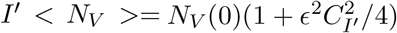. This yields the following expression for *P*:

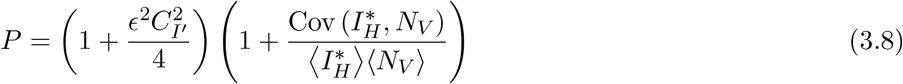

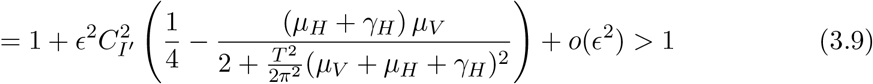

which also yields:

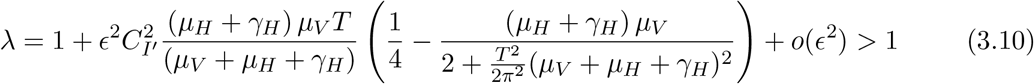

Interestingly the covariance between the fluctuations of the density of infected hosts and the density of the vector population in (3.8) remains negative (as in model *I*, see **Figure S1**). Yet, the effect of seasonality on the average density of the vector population is positive and overwhelms its effect on the covariance. This explains why seasonality has a positive effect on pathogen persistence in model *I*′ (**Figure 2**).

#### 3.2.2 Fluctuations in biting rates

In model *II* we have *C_I_* = 0 and *C_II_* = 1 and equation (3.2) reduces to

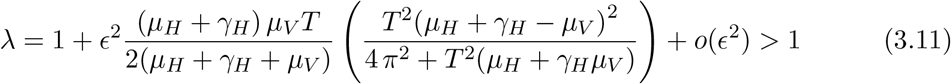

In other words, seasonality is increasing disease persistence in this model (**Figure 2**). Again, we can use equation (2.6) for *P* to get a better understanding of this result. Since seasonality affects only the biting rate *a* the density of the vector population *N_V_* is constant and we get

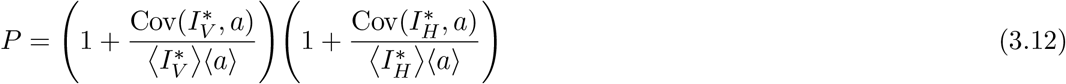

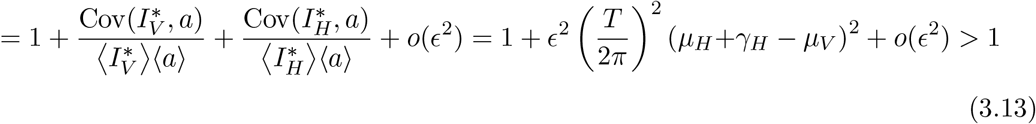

To understand the sign of *P* – 1 we need to understand the sign of the covariances that appear in (3.13). Crucially, we obtained approximations for these covariances in the Supplementary Information, and we show that the sign of 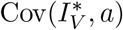 is the sign of (*μ_H_* + *γ_H_*) – *μ_V_* and is the opposite sign of 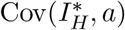. Indeed, the duration of infection in the host, 1/(*μ_H_* + *γ_H_*), and in the vector, 1/*μ_V_*, govern the speed at which the dynamics of the infections reacts to a fluctuation of the biting rate. Shorter duration of infection in one host relative to the other leads to a positive covariance with the biting rate as well as larger amplitude of fluctuations because the epidemic spreads faster with shorter generation time (see **Figure 5**). Interestingly, the sum of these two covariances is always positive, unless the duration of infection in both hosts is equal (i.e., *μ_H_* + *γ_H_* = *μ_V_*). **Figure 5** illustrates the influence of the relative duration of infection in the two hosts on the sign of the covariances and on the amplitude of the oscillations.

**Figure 5:**
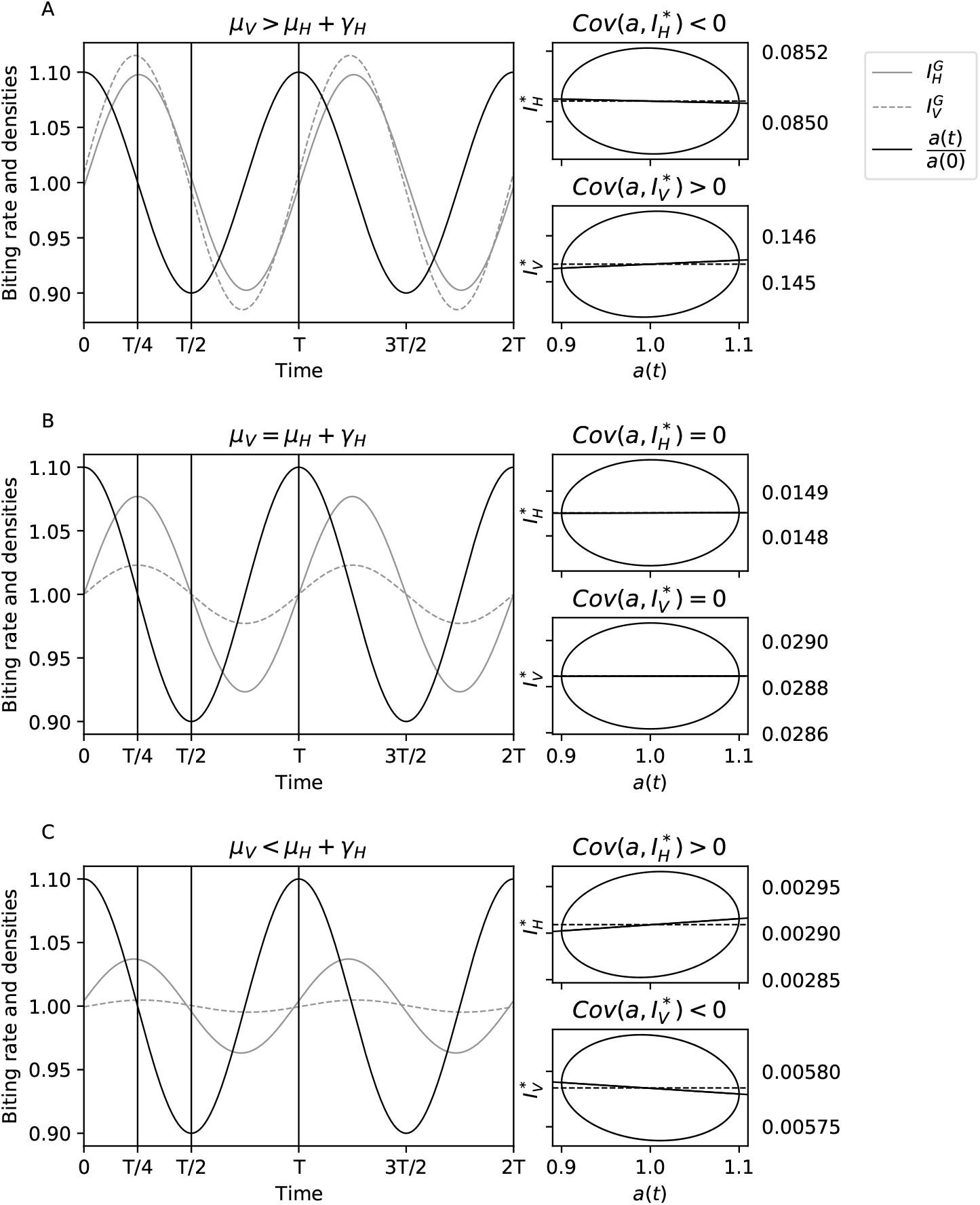
Understanding the covariance between *a*, 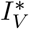 and 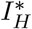 in model *II*. We plot the joint dynamics of the biting rate (solid black line), the density of infected vectors (dashed gray line with 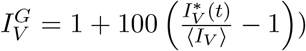 and the density of infected hosts (full gray line with 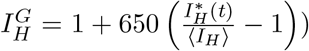 in different scenarios. In **(A**) we assume *γ_H_* = 0.01 and thus *μ_V_* > *μ_H_* + *γ_H_*. In this case the lag between the fluctuation of *a*(*t*) and *I_V_*(*t*)* is lower than *T*/4 which leads to 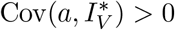 as indicated on the phase diagram on the right panel. In contrast, the lag between the fluctuation of *a*(*t*) and *I_V_*(*t*)* is higher than *T*/4 which leads to 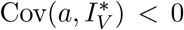. As in Figure 2 the sign of the covariance is given by the slope of the regression line indicated on the phase plane with a black line. In **(B**) we assume *γ_H_* = 0.49 and thus *μ_V_* = *μ_H_* + *γ_H_*. In this case the lag between the fluctuation of *a*(*t*) and both *I_V_*(*t*)* and *I_H_*(*t*)* is equal to *T*/4 which implies that 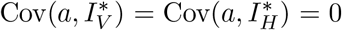. In case **(C**) we assume *γ_H_* = 1.25 which implies that *μ_V_* < *μ_H_* + *γ_H_*. Compare with **(A**) and note how the modification of a single parameter affects the sign of both 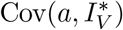 and 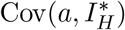. Other parameter values: *ϵ* = 0.5, *α*_0_ = *N_V_*(0) = *β_HV_* = *β_VH_* = 1, *γ_V_* = 0.5, *μ_H_* = 0.01, *T* = 1.

## 4. Discussion

Periodic fluctuations of the environment due to seasonality can affect dramatically the dynamics of infectious diseases [1, 14]. In vector-borne diseases, understanding the overall influence of seasonality is difficult because fluctuations in temperature alter multiple life-history traits of the vector and the pathogen [15, 16, 17, 18, 19]. Hence multiple steps of the life-cycle of the pathogen may be affected simultaneously by seasonality. Previous theoretical analysis of the influence of seasonality focused mainly on the influence of fluctuations in vector density [3, 7, 13] and showed how these fluctuations could either increase or decrease the persistence of vector-borne pathogens. Fluctuations in transmission rates have also been shown to affect the persistence of vector-borne pathogens [20]. Yet, we currently lack a good understanding of the influence of seasonality when the periodic fluctuations of the environment can affect multiple components of the pathogen life-cycle. Here we expand earlier studies of the effect of seasonality to provide a deeper understanding of the influence of periodic fluctuations of the environment on the basic reproduction ratio (and thus on the persistence) of vector-borne pathogens. Pathogen persistence varies with the speed, the amplitude and the shape of the fluctuations as well as on the specific life-history traits modified by seasonality.

### The speed and the amplitude of seasonal fluctuations

Our analysis shows that lower speed of fluctuations (i.e., higher *T*) and higher amplitudes (i.e., higher *ϵ*) always magnify the effects of seasonality. Indeed, faster fluctuations (e.g., daily) or fluctuations with small amplitudes tend to average out and they tend to have a negligible effect on epidemiological dynamics.

### The shape of seasonal fluctuations

If the periodic functions *f*(*t*) used to model the effect of seasonality are such that < *f* >≠ 0 we expect a first order effect of seasonality. This effect simply results from the effect of seasonality on transmission opportunities. For instance, if the influence of seasonality is only to increase the duration of a winter season in which the density of vectors is very low, seasonality will always result in lower disease persistence because it will decrease the average density of vectors. We illustrated this scenario in **Figure 1**. In contrast, if we use periodic functions where < *f* >= 0 this first order effect of seasonality vanishes. Yet, seasonality can still have a second order effect on pathogen persistence. This second order effect can be qualitatively different from the first order effect (compare **Figures 1** and **2**). It is interesting to note that this second order effect can also be affected by the shape of the fluctuations. This is illustrated by the difference between model *I* and model *I*′ (see **Figures 2**). Both models assume that seasonality affect the fluctuation of vector density but the shape of the fluctuation varies between the two models (**4**) and affects qualitatively the influence of seasonality on pathogen persistence.

### The life-history traits altered by seasonal fluctuations

As pointed out before, the influence of seasonality varies with the life-history traits affected by seasonality [3, 8, 9, 20]. In the present paper we focused on the influence of seasonality on two quantities: the vector density (model *I*) and the biting rate of the vector (model *II*). At the first order, fluctuations in both quantities had similar effects on pathogen persistence because both quantities are linked to pathogen transmission. Yet, it is interesting to note that fluctuations in the biting rate where twice more impactful. This factor 2 stems from the fact that biting rates act at two different stages of the pathogen life-cycle, while vector density acts only once. This is clear from the expression of *R*_0_ in equation (2.2) where *α* appears twice and *N_V_* only once.

At the second order, the influence of seasonality varies between models *I* and *II* and when seasonality acts on both the biting rate and the vector density, the overall influence of seasonality depends on the magnitude of the influence of seasonality on these two traits (i.e., the parameters *C_I_* and *C_II_* in equation (3.4)). A better understanding of these effects can be obtained from the examination of *P* and the covariance between different dynamical variables of the system. In a fluctuating environment, several dynamical variables change through time and the phase shift between these quantities determine if the average transmission is increased or decreased by seasonality. Indeed, if *X* and *Y* refer to dynamical quantities involved in transmission, the average transmission will given by < *XY* >=< *X* >< *Y* > +Cov(*X, Y*) and hence by the covariance between these quantities. The analysis of models *I* and *II* illustrate how one can gain an intuitive understanding of the effect of seasonality via the examination of these covariance terms. Our examination of the covariance is akin to the interpretation of the effect of seasonality mediated by the relative timing between the peaks in prevalence in the vector and in the host populations [7, 13].

To conclude, we present a general theoretical framework that allows us to extend previous analyses of the influence of seasonal fluctuations of the environment on the *R*_0_ of vector-borne pathogens. Our analysis highlights the complexity of the influence of seasonality and the necessity to take into account the details of the biology of the pathogen and the vector to understand the effect of seasonality because very similar models can yield qualitatively different conclusions on the influence of seasonality. We hope the present theoretical framework will be used to explore the influence of seasonality in a broader range of epidemiological scenarios tailored to the biology of different infectious diseases. This will help improve the accuracy of risk maps aiming to identify geographic regions that are most likely to be subject to the emergence or the re-emergence of some pathogens [10, 11, 12]. In addition, our analysis could also help identify more effective time-varying control measures against pathogens [4, 6].

## Notes

### Competing Interest Statement

The authors have declared no competing interest.

