## Supplementary Information for "Seasonality and the persistence of vector-borne pathogens"

December 13, 2022

### 1 The vector-borne disease model

We consider the following dynamical system

$$\begin{aligned}\frac{\mathrm{d}S_H}{\mathrm{d}t} &= \theta - a\beta_{VH}\frac{I_V S_H}{N_H} - \mu_H S_H,\\ \frac{\mathrm{d}I_H}{\mathrm{d}t} &= a\beta_{VH}\frac{I_V S_H}{N_H} - (\mu_H+\gamma_H)I_H,\\ \frac{\mathrm{d}R_H}{\mathrm{d}t} &= \gamma_H I_H - \mu_H R_H,\\ \frac{\mathrm{d}S_V}{\mathrm{d}t} &= \rho_V N_V - a\beta_{HV} I_H S_V - \mu_V S_V,\\ \frac{\mathrm{d}I_V}{\mathrm{d}t} &= a\beta_{HV} I_H S_V - \mu_V I_V,\end{aligned} \tag{1.1}$$

For the sake of simplicity we assume that the total density of the human population $S_H + I_H + R_H = N_H$ remains constant which implies that $\theta = \mu_H\,(S_H + I_H + R_H)$.

### 2 The constant case

In a constant environment we assume that the parameters that govern the life cycle of the vector do not vary with time and if the fecundity of the vector compensates exactly its mortality (i.e., $\rho_V = \mu_V$) the total density of the vector population $S_V + I_V = N_V$ is also constant. In this scenario the basic reproduction ratio is readily derived from the linearisation of the above system at the disease free equilibrium ($I_V = I_H = R_H = 0, S_H = N_H, S_V = N_V$) which yields (see [1] ):

$$\frac{d\mathbf{X}}{dt} = \boldsymbol{A}\,\boldsymbol{X}(t)\,, \quad \text{with } \boldsymbol{X}(t) = (I_H(t), I_V(t))^\top\,, \tag{2.1}$$

with

$$\boldsymbol{A} = \begin{pmatrix} -(\mu_H+\gamma_H) & a\beta_{VH} \\ a\beta_{HV}N_V & -\mu_V \end{pmatrix} \tag{2.2}$$

To determine the ability of the pathogen to invade the population. We use one of the three thresholds $P, \lambda$ and $R_0$ was introduced by [2, 3] and [6] (see the 2.4, 2.5 on the main file ).
We rewrite the constant matrix $\boldsymbol{A} = \boldsymbol{F} - \boldsymbol{V}$ such that $\boldsymbol{F}$ and $\boldsymbol{V}$ satisfies the Next Generation Theorem [1] and yields

$$\boldsymbol{F} = \begin{pmatrix} 0 & a\beta_{VH} \\ a\beta_{HV}N_V & 0 \end{pmatrix} \quad \boldsymbol{V} = \begin{pmatrix} \mu_H+\gamma_H & 0 \\ 0 & \mu_V \end{pmatrix}$$

Therefore

$$\boldsymbol{F}\boldsymbol{V}^{-1} = \begin{pmatrix} 0 & \dfrac{a\beta_{VH}}{\mu_V} \\ \dfrac{a\beta_{HV}N_V}{\mu_H+\gamma_H} & 0 \end{pmatrix},$$

and

$$R_0 = \rho\left(\boldsymbol{F}\boldsymbol{V}^{-1}\right) = \frac{a\beta_{VH}}{\mu_V}\frac{a\beta_{HV}N_V}{\mu_H+\gamma_H}\,. \tag{2.3}$$

### 3 The periodic case

We assume that the vector population and the biting rate oscillate using the following periodic functions

$$N_V(t) = N_V(0)(1+\epsilon C_I\ f(t)) \tag{3.1}$$
$$a(t) = a_0(1+\epsilon\, C_{II} f(t))\ , \tag{3.2}$$

where $f(t)$ is a periodic function with period $T$ and $\epsilon$ measures the amplitude of the effects of seasonal fluctuations. Taylor expansion of the transition matrix $\boldsymbol{A}(t)$ for small $\epsilon$ yields:

$$\boldsymbol{A}(t) = \boldsymbol{A}_0 + \epsilon f(t)\boldsymbol{A}_1 + \epsilon^2 f(t)^2\boldsymbol{A}_2 + o(\epsilon^2) \tag{3.3}$$

with $o(\epsilon^2)$ uniform in time and

$$\boldsymbol{A}_0 = \begin{pmatrix} -(\mu_H+\gamma_H) & a_0\beta_{VH} \\ a_0\beta_{HV}N_V(0) & -\mu_V \end{pmatrix} \tag{3.4}$$
$$\boldsymbol{A}_1 = \begin{pmatrix} 0 & a_0\beta_{VH}C_{II} \\ a_0\beta_{HV}N_V(0)(C_I+C_{II}) & 0 \end{pmatrix} \tag{3.5}$$
$$\boldsymbol{A}_2 = \begin{pmatrix} 0 & 0 \\ a_0\beta_{HV}N_V(0)C_IC_{II} & 0 \end{pmatrix}\,. \tag{3.6}$$

We let $\boldsymbol{L} = \boldsymbol{\Phi}_{\mathbf{A}}(T)$ be the monodromy matrix associated with $\boldsymbol{A}(t)$. We first observe that thanks to Duhamel's formula (see subsection 5.2) for perturbation of linear operators we have the Taylor expansion

$$\mathbf{L} = \mathbf{L}_0 + \epsilon\mathbf{L}_1 + \epsilon^2\mathbf{L}_2 + o(\epsilon^2) \tag{3.7}$$

where

$$\mathbf{L}_0 = e^{T\boldsymbol{A}_0}$$
$$\mathbf{L}_1 = \int_0^T e^{(T-s)\boldsymbol{A}_0} f(s)\boldsymbol{A}_1 e^{s\boldsymbol{A}_0}\,ds$$
$$\mathbf{L}_2 = \int_0^T e^{(T-s)\boldsymbol{A}_0} f(s)^2\boldsymbol{A}_2 e^{s\boldsymbol{A}_0}\,ds + \int_0^T e^{(T-s)\boldsymbol{A}_0} f(s)\boldsymbol{A}_1\int_0^s e^{(s-\tau)\boldsymbol{A}_0} f(\tau)\boldsymbol{A}_1 e^{\tau\boldsymbol{A}_0}\,d\tau\, e^{s\boldsymbol{A}_0}\,ds\,.$$

Since $\boldsymbol{A}_0$ is cooperative (with non negative off diagonal entries) and irreducible, the matrix $\mathbf{L}_0$ has positive entries, and by Perron Frobenius theorem, the spectral radius $\lambda_0 = \rho(\mathbf{L}_0)$ is an isolated eigenvalue with positive left and right eigenvectors $\mathbf{v}_0, \mathbf{u}_0$. Therefore, see e.g. Kato [4] or Kloeckner [5], for small $\epsilon$ there exists positive left and right eigenvectors of $\mathbf{L}$ with eigenvalue $\lambda = \rho(\mathbf{L})$ : we have

$$\mathbf{L}\mathbf{u} = \lambda\mathbf{u}\,,\quad \mathbf{v}\mathbf{L} = \lambda\mathbf{v}\,,\quad \mathbf{v}\mathbf{u} = 1 = (1,1)^T\mathbf{u}\,, \tag{3.8}$$

and we have Taylor expansions for $\lambda, \mathbf{u}, \mathbf{v}$ and therefore for $P$:

$$\mathbf{u} = \mathbf{u}_0 + \epsilon\mathbf{u}_1 + \epsilon^2\mathbf{u}_2 + o(\epsilon^2) \tag{3.9}$$
$$\lambda = \lambda_0 + \epsilon\lambda_1 + \epsilon^2\lambda_2 + o(\epsilon^2) \tag{3.10}$$
$$P = P_0 + \epsilon P_1 + \epsilon^2 P_2 + o(\epsilon^2)\,. \tag{3.11}$$

**Assumption** Without loss in generality, and to simplify statements, we shall assume that $R_0 = P_0 = \lambda_0 = 1$. Indeed the transformation $\boldsymbol{A}_0 \to \boldsymbol{A}_0 + \kappa I$ translates to $\lambda \to \lambda e^{\kappa T}$.

#### 3.1 First order results

We shall assume that $\langle f\rangle = \int_0^1 f(s)\,ds \neq 0$.

**Proposition 3.1.** *The first order approximations of $\lambda$ and $P$ are given by:*

$$\lambda = 1 + \epsilon\langle f\rangle T(C_I + 2C_{II})\frac{\mu_V(\mu_H+\gamma_H)}{\mu_V+\mu_H+\gamma_H} + o(\epsilon)\,. \tag{3.12}$$

$$P = 1 + \epsilon\langle f\rangle T(C_I + 2C_{II}) + o(\epsilon)\,. \tag{3.13}$$

*Proof.* Identifying the coefficient of $\epsilon$ in $\mathbf{L}\mathbf{u} = \lambda\mathbf{u}$ yields:

$$\mathbf{L}_1\mathbf{u}_0 + \mathbf{L}_0\mathbf{u}_1 = \lambda_1\mathbf{u}_0 + \lambda_0\mathbf{u}_1\,. \tag{3.14}$$

Multiplying on the left by $\mathbf{v}_0$, we get since $\mathbf{v}_0\mathbf{L}_0 = \lambda_0\mathbf{v}_0$ and $\mathbf{v}_0\mathbf{u}_0 = 1$,

$$\lambda_1 = \mathbf{v}_0\mathbf{L}_1\mathbf{u}_0 = \int_0^T \mathbf{v}_0 e^{(T-s)\boldsymbol{A}_0} f(s)\boldsymbol{A}_1 e^{s\boldsymbol{A}_0}\mathbf{u}_0\,ds\,. \tag{3.15}$$

Since $\boldsymbol{A}_0\mathbf{L}_0 = \boldsymbol{A}_0 e^{T\boldsymbol{A}_0} = e^{T\boldsymbol{A}_0}\boldsymbol{A}_0 = \mathbf{L}_0\mathbf{u}_0$ and $\mathbb{R}\mathbf{u}_0$ is the eigenspace of $\mathbf{L}_0$ with eigenvalue $\mathbf{u}_0$, we infer that $\mathbf{u}_0$ is an eigenvector of $\boldsymbol{A}_0$ and since $e^{T\boldsymbol{A}_0}\mathbf{u}_0 = \lambda_0\mathbf{u}_0 = \mathbf{u}_0$ we get that $\boldsymbol{A}_0\mathbf{u}_0 = 0$. Similarly $\mathbf{v}_0\boldsymbol{A}_0 = 0$ and thus $e^{s\boldsymbol{A}_0}\mathbf{u}_0 = \mathbf{u}_0$ and $\mathbf{v}_0 e^{s\boldsymbol{A}_0} = \mathbf{v}_0$. Eventually, we get

$$\lambda_1 = T\langle f\rangle\mathbf{v}_0\boldsymbol{A}_1\mathbf{u}_0 = T(C_I+2C_{II})\frac{\mu_V(\mu_H+\gamma_H)}{\mu_V+\mu_H+\gamma_H}. \tag{3.16}$$

We have used explicit formulas for $\mathbf{u}_0$ and $\mathbf{v}_0$. Let $\alpha = a_0\beta_{VH}$ and $\gamma = a_0\beta_{HV}N_V(0) = \frac{\mu_V(\mu_H+\gamma_H)}{\alpha}$. Then

$$\mathbf{u}_0 = \frac{1}{\alpha+\mu_H+\gamma_H}(\alpha, \mu_H+\gamma_H)^T\,, \qquad \mathbf{v}_0 = \frac{\alpha+\mu_H+\gamma_H}{\alpha(\mu_H+\gamma_H+\mu_V)}(\mu_V,\alpha)^T \tag{3.17}$$

To obtain the first order expansion of $P$, we recall formula (2.7) of the main text: since for $\epsilon = 0$, we have $P_0 = R_0 = 1$, we have

$$P = P_0\frac{\langle a\rangle\langle aN_V\rangle}{a_0^2N_V(0)}\left(1+\frac{\mathrm{Cov}(a,N_V)}{\langle a\rangle\langle N_V\rangle}\right)\left(1+\frac{\mathrm{Cov}(a,I_V^*)}{\langle a\rangle\langle I_V^*\rangle}\right)\left(1+\frac{\mathrm{Cov}(aN_V,I_H^*)}{\langle aN_V\rangle\langle I_H^*\rangle}\right). \tag{3.18}$$

Recall that we have

$$\mathbf{X}^*(t) = \boldsymbol{\Phi}_{\mathbf{A}}(t)\mathbf{u} = e^{t\boldsymbol{A}_0}\mathbf{u}_0 + o(1) = \mathbf{u}_0 + o(1)\,. \tag{3.19}$$

Therefore, since $I_V^* = e_2^T\mathbf{X}^*$ is the second coordinate of $\mathbf{X}^*$, and the covariance with a constant is 0,

$$\mathrm{Cov}(a, I_V^*) = \mathrm{Cov}(a_0(1+C_{II}\epsilon f(s)), e_2^T\mathbf{X}^*(s)) = a_0C_{II}\epsilon\,\mathrm{Cov}(f(s), e_2^T\mathbf{u}_0) + o(\epsilon) = o(\epsilon)\,. \tag{3.20}$$

Similarly $\mathrm{Cov}(aN_V, I_H^*) = o(\epsilon)$ and $\mathrm{Cov}(a, N_V) = C_IC_{II}\epsilon^2\,\mathrm{Var}(f) = o(\epsilon)$. Hence

$$\begin{aligned}P &= \frac{\langle a\rangle\langle aN_V\rangle}{a_0^2N_V} + o(\epsilon)\\ &= \langle 1+\epsilon C_{II}f(s)\rangle\,\langle(1+\epsilon C_If(s)+o(\epsilon))(1+\epsilon C_{II}f(s))\rangle + o(\epsilon)\\ &= 1+\epsilon(C_I+2C_{II}) + o(\epsilon)\,.\end{aligned}$$

□

#### 3.2 Second order results

##### 3.2.1 Expansion for $\lambda$

**Proposition 3.2.** *With $w = \mu_H+\gamma_H+\mu_V$, $f(t) = \cos(2\pi t/T)$ and $c = 1 + w^2\left(\frac{T}{2\pi}\right)^2$ we have the second order Taylor expansion*

$$\lambda = 1 + \epsilon^2\lambda_2 + o(\epsilon^2)\,, \tag{3.21}$$

*where*

$$\begin{aligned}\lambda_2 &= \boldsymbol{v}_0\mathbf{L}_2\boldsymbol{u}_0 + (1-e^{-Tw})^{-1}\,\boldsymbol{v}_0(\mathbf{L}_1)^2\boldsymbol{u}_0\\ &= \frac{(\mu_H+\gamma_H)\mu_VT}{2(\gamma_H+\mu_H+\mu_V)}\left(C_IC_{II} - \frac{T^2((C_I+C_{II})\mu_V - C_{II}(\mu_H+\gamma_H))((C_I+C_{II})(\mu_H+\gamma_H)-C_{II}\mu_V)}{4\pi^2+T^2(\gamma_H+\mu_H+\mu_V)^2}\right).\end{aligned}$$

*Proof.* Without loss in generality we can assume $\gamma_H = 0$ and then replace in the final formulas $\mu_H$ by $\mu_H + \gamma_H$. Since $\lambda_1 = const.T\langle f\rangle = 0$, we obtain from (3.14)

$$\mathbf{L}_1\mathbf{u}_0 = \left(I - e^{T\boldsymbol{A}_0}\right)\mathbf{u}_1 \tag{3.22}$$

Observe that the spectrum of $\boldsymbol{A}_0$ is $\sigma(\boldsymbol{A}_0) = \{0, -w\}$ where $w = -\text{trace}(\boldsymbol{A}_0)$ Let $\pi(x) = x - \langle x, \mathbf{v}_0\rangle\,\mathbf{u}_0$ be the projection on $\mathbf{v}_0^{\perp}$ of the decomposition $\mathbb{R}^2 = \mathbf{v}_0^{\perp} \oplus \mathbb{R}\mathbf{u}_0$. Since $\mathbf{v}_0\mathbf{L}_1\mathbf{u}_0 = \lambda_1 = 0$ we have

$$\mathbf{u}_1 = \left(I - e^{T\boldsymbol{A}_0}\right)^{-1}|_{G_0}\mathbf{L}_1\mathbf{u}_0\,, \tag{3.23}$$

where $G_0 = {\mathbf{v}_0}^{\perp} = Ker(\boldsymbol{A}_0 + wI)$, $\mathbf{u}_1 \in G_0$. Since if $\mathbf{v}_0 x = 0$ then $\boldsymbol{A}_0 x = -wx$ we obtain

$$\mathbf{u}_1 = \left(1 - e^{-Tw}\right)^{-1}\mathbf{L}_1\mathbf{u}_0 \tag{3.24}$$

Identifying the coefficients of $\epsilon^2$ in $\mathbf{L}_\epsilon = \lambda_\epsilon\mathbf{u}_\epsilon$ one gets

$$\mathbf{L}_2\mathbf{u}_0 + \mathbf{L}_1\mathbf{u}_1 + L_0\mathbf{u}_2 = \lambda_0\mathbf{u}_2 + \lambda_2\mathbf{u}_0$$

Multiplying on the left by $\mathbf{v}_0$, we get, since $\lambda_0 = 1$,$\lambda_1 = 0$, $\mathbf{v}_0\mathbf{u}_0 = 1$, $\mathbf{v}_0\mathbf{L}_0 = \mathbf{v}_0$,

$$\lambda_2 = \mathbf{v}_0\mathbf{L}_2\mathbf{u}_0 + \left(1 - e^{-Tw}\right)^{-1}\mathbf{v}_0\left(\mathbf{L}_1\right)^2\mathbf{u}_0 = \mathbf{v}_0\mathbf{L}_2\mathbf{u}_0 + (1 - e^{-Tw})^{-1}\,\mathbf{v}_0(\mathbf{L}_1)^2\mathbf{u}_0\,. \tag{3.25}$$

From (5.35) and (5.28)

$$\left(1 - e^{-wT}\right)^{-1}\left(\mathbf{L}_1\right)^2\mathbf{u}_0 = \frac{T}{2\pi c}N(T, -w)\left(\boldsymbol{A}_0\boldsymbol{A}_1 - \boldsymbol{A}_1\boldsymbol{A}_0\right)\frac{\boldsymbol{A}_0\boldsymbol{A}_1\mathbf{u}_0}{w^2}.$$

Hence,

$$\begin{aligned}
\lambda_2 &= \mathbf{v}_0\mathbf{L}_2\mathbf{u}_0 + \left(1 - e^{-wT}\right)^{-1}\mathbf{v}_0\left(\mathbf{L}_1\right)^2\mathbf{u}_0 \\
&= -\frac{1}{c}\left(w\left(\frac{T}{2\pi}\right)^2\frac{T}{2} + \frac{T}{2\pi}N(T, -w)\right)\mathbf{v}_0\boldsymbol{A}_1\frac{\boldsymbol{A}_0}{w}\boldsymbol{A}_1\mathbf{u}_0 - \frac{T}{2\pi c}N(T, -w)\mathbf{v}_0\boldsymbol{A}_1\boldsymbol{A}_0\frac{\boldsymbol{A}_0\boldsymbol{A}_1\mathbf{u}_0}{w^2} \\
&\quad + \frac{T}{2}\mathbf{v}_0\left(I + \frac{\boldsymbol{A}_0}{w}\right)\boldsymbol{A}_2\mathbf{u}_0 - \frac{1 - e^{-Tw}}{2c'w^2}\mathbf{v}_0\boldsymbol{A}_0\boldsymbol{A}_2\mathbf{u}_0. \\
&= -\frac{T}{2c}\left(\frac{T}{2\pi}\right)^2\mathbf{v}_0\boldsymbol{A}_1\boldsymbol{A}_0\boldsymbol{A}_1\mathbf{u}_0 + \frac{T}{2}\mathbf{v}_0\left(I + \frac{\boldsymbol{A}_0}{w}\right)\boldsymbol{A}_2\mathbf{u}_0.
\end{aligned}$$

By the definition of $\boldsymbol{A}_0$, $\boldsymbol{A}_1$ and $\boldsymbol{A}_2$, one gets

$$\begin{aligned}
\lambda_2 &= -\frac{T}{2c}\left(\frac{T}{2\pi}\right)^2\mathbf{v}_0\boldsymbol{A}_1\boldsymbol{A}_0\boldsymbol{A}_1\mathbf{u}_0 + \frac{T}{2}\mathbf{v}_0\left(I + \frac{\boldsymbol{A}_0}{w}\right)\boldsymbol{A}_2\mathbf{u}_0. \\
&= -\frac{T}{2c}\left(\frac{T}{2\pi}\right)^2\frac{\mu_V\mu_H}{w}\left(\left(\left(C_I + C_{II}\right)\mu_H - \mu_V C_{II}\right)\left(\left(C_I + C_{II}\right)\mu_V - \mu_H C_{II}\right)\right) + \frac{T}{2}\frac{C_{II}C_I\mu_H\mu_V}{w} \\
&= \frac{T\mu_V\mu_H}{w}\left(\frac{C_{II}C_I}{2} - \frac{1}{2c}\left(\frac{T}{2\pi}\right)^2\left(\left(C_I + C_{II}\right)\mu_H - \mu_V C_{II}\right)\left(\left(C_I + C_{II}\right)\mu_V - \mu_H C_{II}\right)\right)
\end{aligned}$$

□

##### 3.2.2 Expansion for $P$

Let us remember that we assume that $R_0 = \dfrac{a_0^2\beta_{VH}\beta_{HV}N_V(0)}{\mu_V(\mu_H + \gamma_H)} = 1$. Therefore, since $\langle a\rangle = a_0$, $\langle N_V\rangle = N_V(0)$ we get

$$P = \left(1 + \frac{\text{Cov}\,(a, N_V)}{\langle a\rangle\langle N_V\rangle}\right)\left(1 + \frac{\text{Cov}\,(I_V^{\star}, a)}{\langle I_V^{\star}\rangle\langle a\rangle}\right)\left(1 + \frac{\text{Cov}\,(I_H^{\star}, aN_V)}{\langle I_H^{\star}\rangle\langle aN_V\rangle}\right). \tag{3.26}$$

For $f(t) = \cos(2\pi t/T)$ we have $\langle f^2 \rangle = 1/2$. Therefore, the first factor of (3.26) is

$$\frac{\text{Cov}\,(a, N_V)}{\langle a\rangle\langle N_V\rangle} = \text{Cov}(1 + \epsilon C_{II} f, 1 + \epsilon C_I f) = \epsilon^2 C_I C_{II} \langle f^2 \rangle = \frac{\epsilon^2}{2} C_I C_{II}\,. \tag{3.27}$$

The second factor is computed with the help of formula (3.38) of Lemma 3.3 below:

$$\frac{\text{Cov}(I_V^*, a)}{\langle I_V^*\rangle\langle a\rangle} = \frac{\text{Cov}(I_V^*, 1 + \epsilon C_{II} f)}{\langle I_V^*\rangle} \tag{3.28}$$

$$= -\epsilon^2 \frac{C_{II}}{2c} \left(\frac{T}{2\pi}\right)^2 \mu_V \left(-C_{II}\mu_V - C_I\mu_V + C_{II}\mu_H\right) + o(\epsilon) \tag{3.29}$$

Similarly, for the third factor, we get

$$\frac{\text{Cov}\,(I_H^\star, aN_V)}{\langle I_H^\star\rangle\langle aN_V\rangle} = \frac{\text{Cov}(I_H^*, (1 + \epsilon C_{II} f)(1 + \epsilon C_I f))}{\langle I_H^*\rangle\langle (1 + \epsilon C_{II} f)(1 + \epsilon C_I f)\rangle} \tag{3.30}$$

$$= \epsilon\left( (C_I + C_{II}) \frac{\langle I_H^*, f\rangle}{\langle I_H^*\rangle (1 + \frac{1}{2}\epsilon^2 C_I C_{II})} + \epsilon C_I C_{II} \frac{\langle I_H^*, f^2\rangle}{\langle I_H^*\rangle (1 + \frac{1}{2}\epsilon^2 C_I C_{II})}\right) \tag{3.31}$$

$$= -(C_I + C_{II}) \frac{\epsilon^2}{2c} \left(\frac{T}{2\pi}\right)^2 \mu_H(-C_{II}\mu_H + C_I\mu_V + C_{II}\mu_V) + \frac{1}{2}\epsilon^2 C_I C_{II} + o(\epsilon^2)\,. \tag{3.32}$$

Combining all these yields, since $c = 1 + w^2 \left(\frac{T}{2\pi}\right)^2$,

$$\boxed{P = 1 + \epsilon^2 \left[\frac{C_{II} C_I}{2} - \frac{1}{2c}\left(\frac{T}{2\pi}\right)^2 \left((C_I + C_{II})\,\mu_V - C_{II}\mu_H\right)\left((C_I + C_{II})\,\mu_H - C_{II}\mu_V\right)\right] + o(\epsilon^2)}. \tag{3.33}$$

We derive equation (3.5) of the main text by letting $C_{II} = 0$ and $C_I = 1$ (and $\mu_H$ is replaced by $\mu_H + \gamma_H$). Similarly, we derive equation (3.12) of the main text by letting $C_I = 0$ and $C_{II} = 1$.

**Lemma 3.3.** *We have,*

$$\langle \boldsymbol{X}^* \rangle = \boldsymbol{u}_0 + \epsilon \frac{T}{2\pi} (I + \frac{\boldsymbol{A}_0}{w}) \boldsymbol{A}_1 \boldsymbol{u}_0 + o(\epsilon)\,. \tag{3.34}$$

$$\langle \boldsymbol{X}^* f \rangle = -\epsilon \frac{1}{2c} \left(\frac{T}{2\pi}\right)^2 \boldsymbol{A}_0 \boldsymbol{A}_1 \boldsymbol{u}_0 + o(\epsilon)\,. \tag{3.35}$$

$$\left\langle \boldsymbol{X}^* f^2 \right\rangle = \frac{1}{2} \boldsymbol{u}_0 + o(\epsilon)\,. \tag{3.36}$$

*Therefore,*

$$\frac{\langle I_H^* f\rangle}{\langle I_H^*\rangle} = -\frac{\epsilon}{2c} \left(\frac{T}{2\pi}\right)^2 \mu_H(-C_{II}\mu_H + C_I\mu_V + C_{II}\mu_V) + o(\epsilon)\,. \tag{3.37}$$

$$\frac{\langle I_V^* f\rangle}{\langle I_V^*\rangle} = -\epsilon \frac{1}{2c} \left(\frac{T}{2\pi}\right)^2 \mu_V \left(-C_{II}\mu_V - C_I\mu_V + C_{II}\mu_H\right) + o(\epsilon)\,. \tag{3.38}$$

*Proof.* We first give an expansion of $\mathbf{X}^*(t)$ itself in terms of powers of $\epsilon$.

$$\begin{aligned}\mathbf{X}^*(t) &= \mathbf{\Phi}_{\mathbf{A}}(t)\mathbf{u}_\epsilon \\ &= \Big[e^{\boldsymbol{A}_0 t} + \epsilon \int_0^t e^{(t-s)\boldsymbol{A}_0} \boldsymbol{A}_1 f(s) e^{s\boldsymbol{A}_0} ds + \epsilon^2 \left(\int_0^t e^{(t-s)\boldsymbol{A}_0} \boldsymbol{A}_1 f(s) \int_0^s e^{(s-\tau)\boldsymbol{A}_0} \boldsymbol{A}_1 f\,(\tau)\, e^{\tau \boldsymbol{A}_0} d\tau ds \right.\\ &\quad \left. + \int_0^t e^{(t-s)\boldsymbol{A}_0} \boldsymbol{A}_2 f^2(s) e^{s\boldsymbol{A}_0} ds\right) + O(\epsilon^2)\Big] \times \left(\mathbf{u}_0 + \epsilon \mathbf{u}_1 + \epsilon^2 \mathbf{u}_2 + O(\epsilon^3)\right)\end{aligned}$$

$$= e^{\boldsymbol{A}_0 t}\mathbf{u}_0 + \epsilon\left(\int_0^t e^{(t-s)\boldsymbol{A}_0}\boldsymbol{A}_1 f(s)e^{s\boldsymbol{A}_0}ds\mathbf{u}_0 + e^{\boldsymbol{A}_0 t}\mathbf{u}_1\right) + \epsilon^2\left(e^{\boldsymbol{A}_0 t}\mathbf{u}_2 + \int_0^t e^{(t-s)\boldsymbol{A}_0}\boldsymbol{A}_1 f(s)e^{s\boldsymbol{A}_0}ds\mathbf{u}_1\right.$$

$$\left. + \int_0^t e^{(t-s)\boldsymbol{A}_0}\boldsymbol{A}_1 f(s)\int_0^s e^{(s-\tau)\boldsymbol{A}_0}\boldsymbol{A}_1 f(\tau)\, e^{\tau\boldsymbol{A}_0}d\tau ds\mathbf{u}_0 + \int_0^t e^{(t-s)\boldsymbol{A}_0}\boldsymbol{A}_2 f^2(s)e^{s\boldsymbol{A}_0}ds\mathbf{u}_0\right) + o(\epsilon).$$

**Computation of $\langle \mathbf{X}^*, f\rangle$**

$$\int_0^T f(t)\,\mathbf{X}^*(t)dt$$
$$= \int_0^T f(t)e^{\boldsymbol{A}_0 t}\mathbf{u}_0 dt + \epsilon\left(\int_0^T f(t)\int_0^t e^{(t-s)\boldsymbol{A}_0}f(s)\,\boldsymbol{A}_1 e^{s\boldsymbol{A}_0}\mathbf{u}_0 ds dt + \int_0^T f(t)e^{\boldsymbol{A}_0 t}\mathbf{u}_1 dt\right) + o(\epsilon)\,.$$

The first term is easily computed

$$\int_0^T f(t)e^{\boldsymbol{A}_0 t}\mathbf{u}_0 dt = \int_0^T f(t)\mathbf{u}_0 dt = T\langle f\rangle\mathbf{u}_0 = 0\,.$$

The first term of the $\epsilon$ factor is:

$$\int_0^T f(t)\int_0^t e^{(t-s)\boldsymbol{A}_0}f(s)\,\boldsymbol{A}_1 e^{s\boldsymbol{A}_0}\mathbf{u}_0 ds dt$$
$$= \int_0^T f(t)\int_0^t f(s)\left(I + \frac{\boldsymbol{A}_0}{w} - \frac{e^{-w(t-s)}\boldsymbol{A}_0}{w}\right)\boldsymbol{A}_1\mathbf{u}_0 ds dt$$
$$= -\int_0^T f(t)\int_0^t f(s)e^{-w(t-s)}ds dt\frac{\boldsymbol{A}_0\boldsymbol{A}_1\mathbf{u}_0}{w}$$
$$= -\frac{1}{c}\left(\frac{wT}{2}\left(\frac{T}{2\pi}\right)^2 + \frac{T}{2\pi}N(T,-w)\right)\frac{\boldsymbol{A}_0\boldsymbol{A}_1\mathbf{u}_0}{w}.$$

The second term of the $\epsilon$ factor is:

$$\int_0^T f(t)e^{\boldsymbol{A}_0 t}\mathbf{u}_1 dt$$
$$= \int_0^T f(t)\left(I + \frac{\boldsymbol{A}_0}{w} - \frac{e^{-wt}\boldsymbol{A}_0}{w}\right)dt\frac{T}{2\pi c}\frac{\boldsymbol{A}_0\boldsymbol{A}_1\mathbf{u}_0}{w}$$
$$= -N(T,-w)\frac{T}{2\pi c w^2}\boldsymbol{A}_0^2\boldsymbol{A}_1\mathbf{u}_0$$
$$= N(T,-w)\frac{T}{2\pi c w}\boldsymbol{A}_0\boldsymbol{A}_1\mathbf{u}_0 \text{ since } \boldsymbol{A}_0^2 = -w\boldsymbol{A}_0.$$

Hence, the $\epsilon$ factor is

$$-\frac{1}{c}\left(\frac{wT}{2}\left(\frac{T}{2\pi}\right)^2 + \frac{T}{2\pi}N(T,-w)\right)\frac{\boldsymbol{A}_0\boldsymbol{A}_1\mathbf{u}_0}{w} + \left(\frac{T}{2\pi}\right)^2\left(\frac{3N(T,-w)}{4cc'} + \frac{wT^2}{4\pi c}\right)\frac{\boldsymbol{A}_1\boldsymbol{A}_0}{w}\boldsymbol{A}_1\mathbf{u}_0 = -\frac{T}{2c}\left(\frac{T}{2\pi}\right)^2\boldsymbol{A}_0\boldsymbol{A}_1\mathbf{u}_0,$$

from which we deduce (3.35).

**Computation of $\langle \mathbf{X}^*\rangle$**

$$\int_0^T \mathbf{X}^*(t)dt = \int_0^T e^{\boldsymbol{A}_0 t}\mathbf{u}_0 dt + \epsilon\left(\int_0^T\int_0^t e^{(t-s)\boldsymbol{A}_0}\boldsymbol{A}_1 f(s)e^{s\boldsymbol{A}_0}\mathbf{u}_0 ds dt + \int_0^T e^{\boldsymbol{A}_0 t}\mathbf{u}_1 dt\right) + o(\epsilon)\,.$$

The first term is

$$\int_0^T e^{\boldsymbol{A}_0 t}\mathbf{u}_0 dt = \int_0^T \mathbf{u}_0 dt = T\mathbf{u}_0$$

From (5.3) and (5.26) we get:

$$\begin{aligned}\int_0^T e^{\boldsymbol{A}_0 t}\mathbf{u}_1 dt &= (1-e^{-Tw})^{-1}\frac{N_0(T,-w)}{w}\left(\int_0^T e^{t\boldsymbol{A}_0}\boldsymbol{A}_0\,dt\right)\boldsymbol{A}_1\mathbf{u}_0\\ &= (1-e^{-Tw})^{-1}\frac{N_0(T,-w)}{w}(e^{T\boldsymbol{A}_0}-I)\boldsymbol{A}_1\mathbf{u}_0\\ &= N(T,-w)\frac{\boldsymbol{A}_0\boldsymbol{A}_1\mathbf{u}_0}{w^2}\,.\end{aligned}$$

Eventually,

$$\begin{aligned}&\int_0^T\int_0^t e^{(t-s)\boldsymbol{A}_0}\boldsymbol{A}_1 f(s)e^{s\boldsymbol{A}_0}dsdt\mathbf{u}_0\\ &=\frac{T^2}{2\pi}\left(I+\frac{\boldsymbol{A}_0}{w}\right)\boldsymbol{A}_1\left(I+\frac{\boldsymbol{A}_0}{w}\right)\mathbf{u}_0-\frac{1}{c}\left(-2w\left(\frac{T}{2\pi}\right)^2N(T,-w)+\frac{T^2}{2\pi}\right)\left(I+\frac{\boldsymbol{A}_0}{w}\right)\boldsymbol{A}_1\frac{\boldsymbol{A}_0\mathbf{u}_0}{w}\\ &\quad-\frac{N(T,-w)}{w}\frac{\boldsymbol{A}_0}{w}\boldsymbol{A}_1\left(I+\frac{\boldsymbol{A}_0}{w}\right)\mathbf{u}_0+\frac{N(T,-w)}{w}\frac{\boldsymbol{A}_0\boldsymbol{A}_1\boldsymbol{A}_0\mathbf{u}_0}{w^2}.\end{aligned}$$

Combining the preceding we get (3.34).
**Computation of $\langle \mathbf{X}^* f^2\rangle$** Since $e^{t\boldsymbol{A}_0}\mathbf{u}_0=\mathbf{u}_0$, we get

$$\int_0^T f^2(t)\mathbf{X}^*(t)dt=\int_0^T \sin\left(\frac{2\pi t}{T}\right)^2 e^{\boldsymbol{A}_0 t}\mathbf{u}_0 dt+o(\epsilon)=\frac{T}{2}\mathbf{u}_0+o(\epsilon)\,. \tag{3.39}$$

**Computation of $\frac{\langle I_H^* f\rangle}{\langle I_H^*\rangle}$** if $e_1=(1,0), e_2=(0,1)$ are the coordinate vectors, then $I_H^*=e_1^T\mathbf{X}^*$. Therefore, $\langle I_H^* f\rangle = e_1^T\langle \mathbf{X}^* f\rangle$ and thanks to the matrix computations

$$\begin{aligned}e_1^T\boldsymbol{A}_0\boldsymbol{A}_1\mathbf{u}_0 &= \frac{\alpha\mu_H\left(-C_{II}\mu_H+C_I\mu_V+C_{II}\mu_V\right)}{\alpha+\mu_H}\\ e_2^T\boldsymbol{A}_0\boldsymbol{A}_1\mathbf{u}_0 &= \frac{\mu_H\mu_V\left(-C_{II}\mu_V-C_I\mu_V+C_{II}\mu_H\right)}{\alpha+\mu_H}\,,\end{aligned} \tag{3.40}$$

we get

$$\begin{aligned}\frac{\langle I_H^* f\rangle}{\langle I_H^*\rangle} &= \frac{\int_0^T f(t)e_1^T\mathbf{X}^*(t)dt}{\int_0^T e_1^T\mathbf{X}^*(t)dt}\\ &= \left(-\epsilon\frac{T}{2c}\left(\frac{T}{2\pi}\right)^2 e_1^T\boldsymbol{A}_0\boldsymbol{A}_1\mathbf{u}_0+o(\epsilon)\right)\frac{1}{Te_1^T\mathbf{u}_0}\left(1-\epsilon\frac{T^2}{2\pi}e_1^T\left(I+\frac{\boldsymbol{A}_0}{w}\right)\frac{\boldsymbol{A}_1\mathbf{u}_0}{Te_1\mathbf{u}_0}+o(\epsilon)\right)\\ &= \frac{1}{Te_1^T\mathbf{u}_0}\left(-\epsilon\frac{T}{2c}\left(\frac{T}{2\pi}\right)^2 e_1^T\boldsymbol{A}_0\boldsymbol{A}_1\mathbf{u}_0+o(\epsilon)\right)\\ &= \frac{1}{T\dfrac{\alpha}{\alpha+\mu_H}}\,\frac{-\epsilon T}{2c}\left(\frac{T}{2\pi}\right)^2\frac{\alpha\mu_H\left(-C_{II}\mu_H+C_I\mu_V+C_{II}\mu_V\right)}{\alpha+\mu_H}+o(\epsilon)\\ &= -\epsilon\frac{1}{2c}\left(\frac{T}{2\pi}\right)^2\mu_H\left(-C_{II}\mu_H+C_I\mu_V+C_{II}\mu_V\right)+O(\epsilon)\end{aligned} \tag{3.41}$$

Similarly,

$$\begin{aligned}\frac{\langle I_V^* f\rangle}{\langle I_V^*\rangle} &= \frac{\int_0^T f(t)e_2^T x(t)dt}{\int_0^T e_2^T x(t)dt}\\ &= \left(-\epsilon\frac{T}{2c}\left(\frac{T}{2\pi}\right)^2 e_2^T\boldsymbol{A}_0\boldsymbol{A}_1\mathbf{u}_0+O(\epsilon)\right)\frac{1}{Te_2^T\mathbf{u}_0}\left(1-\epsilon\frac{T^2}{2\pi}e_2^T\left(I+\frac{\boldsymbol{A}_0}{w}\right)\frac{\boldsymbol{A}_1\mathbf{u}_0}{Te_2^T\mathbf{u}_0}+O(\epsilon)\right)\end{aligned}$$

$$= \frac{1}{Te_2^T \mathbf{u}_0} \left( \frac{-\epsilon T}{2c} \left( \frac{T}{2\pi} \right)^2 e_2^T \boldsymbol{A}_0 \boldsymbol{A}_1 \mathbf{u}_0 \right) + O(\epsilon)$$
$$= \frac{1}{T \frac{\mu_H}{\alpha + \mu_H}} \left[ \frac{-\epsilon T}{2c} \left( \frac{T}{2\pi} \right)^2 \frac{\mu_V \mu_H \left( -C_{II}\mu_V - C_I \mu_V + C_{II}\mu_H \right)}{\alpha + \mu_H} \right] + O(\epsilon)$$
$$= -\epsilon \frac{1}{2c} \left( \frac{T}{2\pi} \right)^2 \mu_V \left( -C_{II}\mu_V - C_I \mu_V + C_{II}\mu_H \right) + O(\epsilon) \tag{3.42}$$

□

### 4 An alternative model for the fluctuation of vector density: Model $I'$

In model I' we assume that $N_V(t) = N_V(0)e^{\epsilon C_{I'} f(t)}$. This results from an alternative model of vector reproduction used in Heesterbeek and Roberts [2, 3]. We assume that the per capita growth rate varies around its mean following

$$\rho_V(t) = \mu_V(1 - \epsilon\kappa \sin(2\pi t/T)) \, . \tag{4.1}$$

Then the total vector population $N_V$ satisfies the ODE

$$\frac{dN_V}{dt} = (\rho_V - \mu_v)N_V, \tag{4.2}$$

and is therefore given by

$$N_V(t) = N_V(0) \exp(\epsilon\kappa \frac{T}{2\pi} \cos(2\pi t/T)) \, , \tag{4.3}$$

that is we take $C_I' = \kappa \frac{T}{2\pi}$.
This yields

$$\boldsymbol{A}_1 = \begin{pmatrix} 0 & 0 \\ a_0 \beta_{HV} N_V(0) C_{I'} & 0 \end{pmatrix} , \qquad \boldsymbol{A}_2 = \begin{pmatrix} 0 & 0 \\ a_0 \beta_{HV} N_V(0) \frac{1}{2} C_{I'}^2 & 0 \end{pmatrix} . \tag{4.4}$$

There is no fluctuation in the biting rate so $\langle a \rangle = a_0$. On the other hand we have $\langle N_V \rangle = N_V(0)(1 + \epsilon^2 \frac{C_{I'}}{4} + o(\epsilon^2))$. The expression of $P$ is thus:

$$P = \frac{\langle N_V \rangle}{N_V(0)} \left( 1 + \frac{\mathrm{Cov}\,(a, N_V)}{\langle a \rangle \langle N_V \rangle} \right) \left( 1 + \frac{\mathrm{Cov}\,(I_V^\star, a)}{\langle I_V^\star \rangle \langle a \rangle} \right) \left( 1 + \frac{\mathrm{Cov}\,(I_H^\star, aN_V)}{\langle I_H^\star \rangle \langle aN_V \rangle} \right) . \tag{4.5}$$
$$= (1 + \epsilon^2 \frac{C_{I'}}{4} + o(\epsilon^2)) \left( 1 + \frac{\mathrm{Cov}\,(I_H^\star, N_V)}{\langle I_H^\star \rangle \langle N_V \rangle} \right) . \tag{4.6}$$

We observe that the second order term, in $\epsilon^2$, of $N_V(t)$ may only gives a term in $\epsilon^3$ in the covariance. In a nutshell, we can keep the same expansion as in the preceding section, the one given by Lemma 3.3, replacing $C_{I'}$ by $C_I$ and $C_{II}$ by 0. Therefore:

$$P = (1 + \epsilon^2 \frac{C_{I'}}{4} + o(\epsilon^2)) \left( 1 - C_{I'} \frac{\epsilon^2}{2c} \left( \frac{T}{2\pi} \right)^2 \mu_V \mu_H + o(\epsilon^2) \right) . \tag{4.7}$$

This leads easily to the expression (3.8) of the main text (if we substitute at the end $\mu_H + \gamma_H$ to $\mu_H$). The fact that $P_\epsilon > 1$ comes from the following sequence of inequalities

$$P = 1 + \frac{\epsilon^2 C_{I'}^2}{2} \left( \frac{1}{2} - \frac{\left(\frac{T}{2\pi}\right)^2 \mu_H \mu_V}{1 + (\mu_H + \mu_V)^2 \left(\frac{T}{2\pi}\right)^2} \right) + o(\epsilon^2)$$
$$\geq 1 + \frac{\epsilon^2 C_{I'}^2}{2} \left( \frac{1}{2} - \frac{1}{4} \frac{\left(\frac{T}{2\pi}\right)^2 (\mu_H + \mu_V)^2}{1 + (\mu_H + \mu_V)^2 \left(\frac{T}{2\pi}\right)^2} \right) + O(\epsilon^2)$$

$$\geq 1 + \frac{\epsilon^2 C_{I'}^2}{8} \frac{\left(\frac{T}{2\pi}\right)^2 (\mu_H + \mu_V)^2}{1 + (\mu_H + \mu_V)^2 \left(\frac{T}{2\pi}\right)^2} + o(\epsilon^2) > 1\,.$$

To compute the influence on $\lambda$ of the model $I'$, we only need to observe that we only need to modify the Taylor expansion(3.3) of $\mathbf{A}(t)$ : we still have no first order term $\lambda = 1 + \epsilon^2 \lambda_2$, and we just plug in the new expressions of $\boldsymbol{A}_1$ and $\boldsymbol{A}_2$ into the formula

$$\lambda_2 = -\frac{T}{2c} \left(\frac{T}{2\pi}\right)^2 \mathbf{v}_0 \boldsymbol{A}_1 \boldsymbol{A}_0 \boldsymbol{A}_1 \mathbf{u}_0 + \frac{T}{2} \mathbf{v}_0 \left(I + \frac{\boldsymbol{A}_0}{w}\right) \boldsymbol{A}_2 \mathbf{u}_0\,,$$

to obtain formula (3.9) of the main text.

### 5 Auxiliary results

#### 5.1 Bounding the flow of an ODE

First we show that control on the coefficients of an ode gives control on the flow of the ode.

**Lemma 5.1.** *Let $\mathbf{A} : [0, a] \to \mathcal{M}_{d\times d}$ be a continuous matrix function, and $\boldsymbol{\Phi}_{\mathbf{A}}(t)$ be its fundamental matrix*

$$\frac{d\boldsymbol{\Phi}_{\mathbf{A}}(t)}{dt} = \mathbf{A}(t)\, \boldsymbol{\Phi}_{\mathbf{A}}(t)\,, \qquad \boldsymbol{\Phi}_{\mathbf{A}}(0) = \mathbf{I}_d\,. \tag{5.1}$$

*Then,*

$$\|\boldsymbol{\Phi}_{\mathbf{A}}(t)\| \leq \exp\left(t \sup_{s\leq t} \|\mathbf{A}(s)\|\right). \tag{5.2}$$

*Proof.* Let $x \in \mathbb{R}^d$. Then $x(t) = \boldsymbol{\Phi}_{\mathbf{A}}(t)x$ is the solution of

$$\frac{dx}{dt} = \mathbf{A}(t)\, x(t)\,, \quad x(0) = x\,. \tag{5.3}$$

Hence,

$$x(t) = x + \int_0^t \mathbf{A}(s)x(s)\, ds, \tag{5.4}$$

and taking norms yield

$$\|x(t)\| \leq \|x\| + \int_0^t \|\mathbf{A}(s)\|\|x(s)\|\, ds \leq \|x\| + \sup_{s\leq t} \|\mathbf{A}(s)\| \int_0^t \|x(s)\|\, ds\,. \tag{5.5}$$

We conclude by Gronwall's Lemma. □

#### 5.2 Duhamel's formula and Taylor expansions

Let us denote by $\boldsymbol{\Phi}_{\mathbf{A}}$ the fundamental matrix of the differential flow generated by the continuous matrix function $t \to \mathbf{A}(t)$. It is the solution of the differential equation

$$\frac{d\boldsymbol{\Phi}_{\mathbf{A}}}{dt} = \mathbf{A}\boldsymbol{\Phi}_{\mathbf{A}}(t) \tag{5.6}$$

with initial condition $\boldsymbol{\Phi}_{\mathbf{A}}(0) = \mathbf{I}$ (the identity matrix).
If $\mathbf{B}(t)$ is another continuous matrix function, then we have

$$\boldsymbol{\Phi}_{\mathbf{A}+\mathbf{B}}(t) = \boldsymbol{\Phi}_{\mathbf{A}}(t)\left(\mathbf{I} + \int_0^t \boldsymbol{\Phi}_{\mathbf{A}}(s)^{-1}\mathbf{B}(s)\, \boldsymbol{\Phi}_{\mathbf{A}+\mathbf{B}}(s) ds\right) \tag{5.7}$$

Let us see how it allows to determine Taylor's expansion. Let us start from

$$\mathbf{A}(t) = \mathbf{A}_0 + \epsilon f(t)\mathbf{A}_1 + o(\epsilon) \tag{5.8}$$

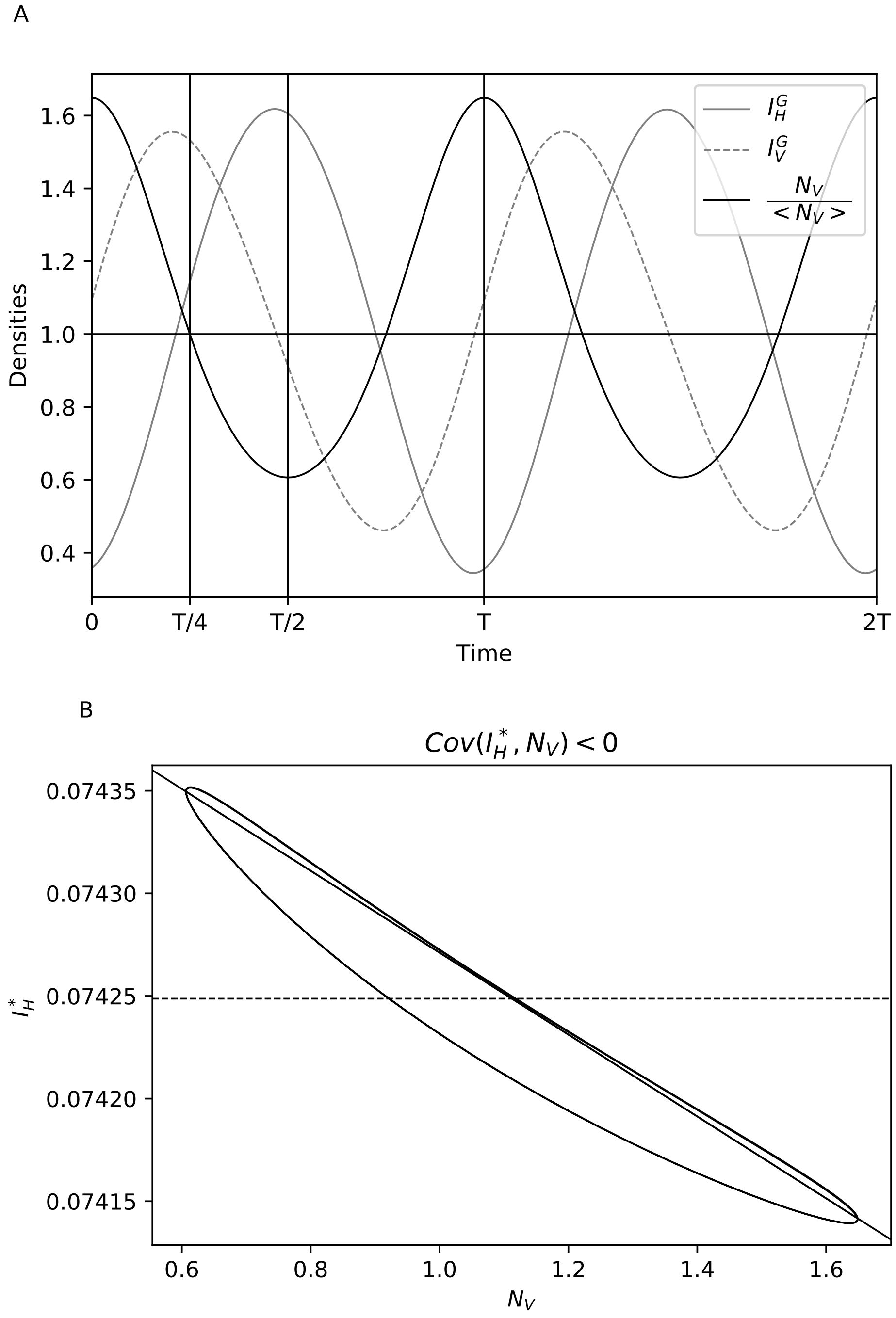


Figure S 1: **Understanding the covariance between $N_V$ and $I_H$ in model $I'$.** The same information as in figure (3) with the infected vector $I_V^G = 1 + 150\left(\frac{I_V(t)}{\langle I_V \rangle} - 1\right)$ and infected host $I_H^G = 1 + 6000\left(\frac{I_H(t)}{\langle I_H \rangle} - 1\right)$. Parameters values: $a_0 = N_V(0) = \beta_{HV} = 1$, $1 = \beta_{VH}, \epsilon = 0.5, T = 1$, $\mu_V = 1, \mu_H = 0.01, \gamma_H = 0.1$.

with $o(\epsilon)$ uniform in time. Then since $\mathbf{\Phi}_{\mathbf{A}_0}(t) = e^{t\mathbf{A}_0}$, by Duhamel's formula

$$\mathbf{\Phi}_{\mathbf{A}}(t) = e^{t\mathbf{A}_0}\left(\mathbf{I} + \int_0^t e^{-s\mathbf{A}_0}(\epsilon f(s)\mathbf{A}_1 + o(\epsilon))\,\mathbf{\Phi}_{\mathbf{A}}(s)\,ds\right) \tag{5.9}$$

Since, by Lemma 5.1, $\sup_{s\leq T}\|\mathbf{\Phi}_{\mathbf{A}}(s)\| \leq C < +\infty$, we have

$$\mathbf{\Phi}_{\mathbf{A}}(t) = e^{t\mathbf{A}_0} + \epsilon\, e^{t\mathbf{A}_0}\int_0^t e^{-s\mathbf{A}_0} f(s)\mathbf{A}_1\,\phi_\epsilon(s)\,ds + o(\epsilon) \tag{5.10}$$

This implies that $\mathbf{\Phi}_{\mathbf{A}}(t) = e^{t\mathbf{A}_0} + o(1)$ which we reinject in the integral term of (5.10) to get

$$\mathbf{\Phi}_{\mathbf{A}}(t) = e^{t\mathbf{A}_0} + \epsilon\, e^{t\mathbf{A}_0}\int_0^t e^{-s\mathbf{A}_0} f(s)\mathbf{A}_1\,e^{s\mathbf{A}_0}\,ds + o(\epsilon) \tag{5.11}$$

Similarly, we obtain the Taylor expansion (3.7).

#### 5.3 Some results of integral calculations.

Recall that $c := 1 + w^2\left(\frac{T}{2\pi}\right)^2$. We let $c' := 1 + w^2\left(\frac{T}{4\pi}\right)^2 = \frac{c+3}{4}$

**Lemma 5.2.** *Let $N(t,x) = \int_0^t f(s)e^{xs}ds$, where $x$ is a real number. Then, for $T, w > 0$,*

$$N(T,w) = \frac{-\frac{T}{2\pi}(e^{wT}-1)}{1+w^2(\frac{T}{2\pi})^2} < 0, \quad N(T,-w) = \frac{-\frac{T}{2\pi}(e^{-wT}-1)}{1+w^2(\frac{T}{2\pi})^2} = -N(T,w)e^{-wT} > 0. \tag{5.12}$$

*Proof.*

$$\begin{aligned}
N(t,k) &= \int_0^t f(s)e^{ks}ds \\
&= \int_0^t \sin(2\pi s/T)e^{ks}ds \\
&= -\frac{T}{2\pi}\int_0^t e^{ks}d\cos(2\pi s/T) \\
&= -\frac{T}{2\pi}\left((e^{kt}\cos(2\pi t/T+) - 1) - k\int_0^t e^{ks}\cos(2\pi s/T+)ds\right) \\
&= -\frac{T}{2\pi}(e^{kt}\cos(2\pi t/T) - 1) + k\left(\frac{T}{2\pi}\right)^2\int_0^t e^{ks}d\sin(2\pi s/T) \\
&= -\frac{T}{2\pi}(e^{kt}\cos(2\pi t/T) - 1) + k\left(\frac{T}{2\pi}\right)^2 e^{kt}\sin(2\pi t/T) - \left(k^2\frac{T}{2\pi}^2\right)N(t,k)
\end{aligned}$$

Therefore,

$$N(t,k) = \frac{-\frac{T}{2\pi}(e^{kt}\cos(2\pi t/T)-1) + k\left(\frac{T}{2\pi}\right)^2(e^{kt}\sin(2\pi t/T))}{1+k^2(\frac{T}{2\pi})^2} \tag{5.13}$$

□

Let us list now some computations that can be established in the same way.

$$\begin{aligned}
\int_0^T f(2s)e^{ws}ds &= \frac{-\frac{T}{4\pi}\left(e^{wT}-1\right)}{c'} = \frac{c}{2c'}N_0(T,w) \\
\int_0^T f(2s)e^{-ws}ds &= \frac{c}{2c'}N_0(T,-w)
\end{aligned}$$

By using integration by parts, one also gets

$$N_3(h,t,k) = \int_0^t \cos(h2\pi s/T)e^{ks}ds = \frac{\frac{T}{2\pi h}e^{kt}\sin\left(2\pi th/T\right) + k\left(\frac{T}{2\pi h}\right)^2\left(e^{kt}\cos\left(2\pi ht/T\right)-1\right)}{1+k^2\left(\frac{T}{2\pi h}\right)^2}.$$

We also list some results in the calculation of the integral.

$$\int_0^T \sin\left(2\pi t/T\right)\int_0^t \sin\left(2\pi s/T\right)\int_0^s \sin\left(2\pi\tau/T\right)d\tau ds dt = 0 \tag{5.14}$$

$$\int_0^T e^{-wt}\sin\left(2\pi t/T\right)\int_0^t \sin\left(2\pi s/T\right)e^{ws}\int_0^s \sin\left(2\pi\tau/T\right)d\tau ds dt = \left(\frac{T}{2\pi}\right)^2\left(\frac{3N(T,-w)}{4cc'}+\frac{wT^2}{4\pi c}\right) \tag{5.15}$$

$$\int_0^T f(t)\int_0^t f(s)\int_0^s f(\tau)e^{-w(s-\tau)}d\tau ds dt = -\frac{1}{c}\left(\frac{T}{4\pi}\right)^2\left(\frac{3N_2(1,T,-w)}{c'}+\frac{wT^2}{\pi}\right) \tag{5.16}$$

$$\int_0^T f(t)e^{-wt}\int_0^t f(s)\int_0^s f(\tau)e^{w\tau}d\tau ds = 0 \tag{5.17}$$

$$\int_0^t f(s)\int_0^s f(\tau)e^{-w\tau}d\tau ds = \frac{5}{3}\frac{N(T,-w)}{c'c''}\left(\frac{T}{4\pi}\right)^2 \tag{5.18}$$

$$\int_0^T f(t)e^{-wt}\int_0^t f(s)e^{ws}\int_0^s f(\tau)e^{-w\tau}d\tau ds dt = \frac{1}{c^2}\left(\frac{T}{4\pi}\right)^2\left(\frac{-8\left(5c-8\right)}{9c''}N(T,-w)+4wT\frac{T}{4\pi}\left(1+e^{-wT}\right)\right) \tag{5.19}$$

$$\int_0^T f(t)\int_0^t f(s)e^{-ws}\int_0^s f(\tau)d\tau ds dt = \left(\frac{T}{2\pi}\right)^2\frac{5N(T,-w)}{6c'c''} \tag{5.20}$$

$$\int_0^T e^{-wt}\sin\left(2\pi t/T\right)\int_0^t \sin\left(2\pi s/T\right)\int_0^s \sin\left(2\pi\tau/T\right)d\tau ds dt = \left(\frac{T}{2\pi}\right)^2\frac{5N(T,-w)}{12c'c''} \tag{5.21}$$

$$\int_0^t \sin(2\pi s/T)e^{ws}\int_0^s \sin(2\pi\tau/T)d\tau ds = -\left(\frac{T}{2\pi}\right)\left(\frac{N_2(2,w,t)}{2}-N_2(1,w,t)\right) \tag{5.22}$$

$$\int_0^t \sin\left(2\pi s/T\right)e^{ws}\int_0^s \sin\left(2\pi\tau/T\right)e^{-w\tau}d\tau ds = \frac{1}{c}\left(\left(\frac{T}{4\pi}\right)^2\left(\cos\left(4\pi\frac{t}{T}\right)-1\right)+\frac{T}{2\pi}N(t,w)\right) \tag{5.23}$$

$$-\frac{w}{2c}\left(\frac{T}{2\pi}\right)^2\left(t-\frac{T}{4\pi}\sin\left(4\pi\frac{t}{T}\right)\right) \tag{5.24}$$

$$\begin{aligned}
&\textstyle\int_0^T\int_0^t \sin 2\pi\frac{s}{T}dsdt = \frac{T^2}{2\pi}\\
&\textstyle\int_0^T\int_0^t \sin 2\pi\frac{s}{T}e^{-ws}dsdt = \frac{1}{c}\left(-2w\left(\frac{T}{2\pi}\right)^2N(T,-w)+\frac{T^2}{2\pi}\right)\\
&\textstyle\int_0^T\int_0^t sin2\pi\frac{s}{T}e^{-w(t-s)}dsdt = \frac{N(T,-w)}{w}\\
&\textstyle\int_0^T e^{-wt}\int_0^t sin2\pi\frac{s}{T}dsdt = \frac{N(T,-w)}{w}
\end{aligned} \tag{5.25}$$

#### 5.4 Computation of $\mathbf{L}_1$ and $\mathbf{L}_2$

Since $\boldsymbol{A}_0$ has two different eigenvalues, 0 and $-w = tr(A)$ we have the formula

$$e^{s\mathbf{A}_0} = I - \frac{1}{w}\left(e^{-ws}-1\right)\boldsymbol{A}_0 \qquad (s\in\mathbb{R}). \tag{5.26}$$

**Lemma 5.3.** *We have*

$$\mathbf{L}_1 = N(T,-w)\frac{(\boldsymbol{A}_0\boldsymbol{A}_1 - \boldsymbol{A}_1\boldsymbol{A}_0)}{w}\,. \tag{5.27}$$

*Since* $\boldsymbol{A}_0\boldsymbol{u}_0 = 0$ *this yields*

$$\mathbf{L}_1\boldsymbol{u}_0 = N_0(T,-w)\frac{\boldsymbol{A}_0\boldsymbol{A}_1\boldsymbol{u}_0}{w}. \tag{5.28}$$

*And thus,*

$$\boldsymbol{u}_1 = (1-e^{-Tw})^{-1}\mathbf{L}_1\boldsymbol{u}_0 = (1-e^{-Tw})^{-1}N_0(T,-w)\frac{\boldsymbol{A}_0\boldsymbol{A}_1\boldsymbol{u}_0}{w}\,. \tag{5.29}$$

*Proof.*

$$\mathbf{L}_1 = \int_0^t e^{(t-s)\boldsymbol{A}_0} f(s)\boldsymbol{A}_1 e^{s\boldsymbol{A}_0} ds \tag{5.30}$$

$$= \int_0^t f(s)\left(\left(I+\frac{\boldsymbol{A}_0}{w}\right) - e^{-w(t-s)}\frac{\boldsymbol{A}_0}{w}\right)\boldsymbol{A}_1\left(\left(I+\frac{\boldsymbol{A}_0}{w}\right) - e^{-ws}\frac{\boldsymbol{A}_0}{w}\right)ds \tag{5.31}$$

$$= \int_0^t f(s)\left[\left(I+\frac{\boldsymbol{A}_0}{w}\right)\boldsymbol{A}_1\left(I+\frac{\boldsymbol{A}_0}{w}\right) - e^{ws}e^{-wt}\frac{\boldsymbol{A}_0}{w}\boldsymbol{A}_1\left(I+\frac{\boldsymbol{A}_0}{w}\right)\right. \tag{5.32}$$

$$\left. -e^{-ws}\left(I+\frac{\boldsymbol{A}_0}{w}\right)\boldsymbol{A}_1\frac{\boldsymbol{A}_0}{w} + e^{-wt}\frac{\boldsymbol{A}_0\boldsymbol{A}_1\boldsymbol{A}_0}{w^2}\right]ds \tag{5.33}$$

$$\Rightarrow \mathbf{L}_1 = \int_0^T f(s)\left[-e^{ws}e^{-wT}\frac{\boldsymbol{A}_0}{w}\boldsymbol{A}_1\left(I+\frac{\boldsymbol{A}_0}{w}\right) - e^{-ws}\left(I+\frac{\boldsymbol{A}_0}{w}\right)\boldsymbol{A}_1\frac{\boldsymbol{A}_0}{w}\right]ds \tag{5.34}$$

$$\mathbf{L}_1 = -N(T,-w)\left(I+\frac{\boldsymbol{A}_0}{w}\right)\boldsymbol{A}_1\frac{\boldsymbol{A}_0}{w} - N(T,w)e^{-wT}\frac{\boldsymbol{A}_0}{w}\boldsymbol{A}_1\left(I+\frac{\boldsymbol{A}_0}{w}\right)$$

$$= N(T,-w)\frac{(\boldsymbol{A}_0\boldsymbol{A}_1 - \boldsymbol{A}_1\boldsymbol{A}_0)}{w}. \tag{5.35}$$

□

**Lemma 5.4.** *We have*

$$\mathbf{L}_2\boldsymbol{u}_0 = \frac{3T}{8\pi c'}N(T,-w)\frac{\boldsymbol{A}_0\boldsymbol{A}_1}{w}\left(I+\frac{\boldsymbol{A}_0}{w}\right)\boldsymbol{A}_1\boldsymbol{u}_0 - \frac{1}{c}\left(w\left(\frac{T}{2\pi}\right)^2\frac{T}{2} + \frac{T}{2\pi}N(T,-w)\right)\left(I+\frac{\boldsymbol{A}_0}{w}\right)\boldsymbol{A}_1\frac{\boldsymbol{A}_0}{w}\boldsymbol{A}_1\boldsymbol{u}_0$$
$$+\frac{3T}{8\pi c'}N(T,-w)\frac{\boldsymbol{A}_0}{w}\boldsymbol{A}_1\frac{\boldsymbol{A}_0}{w}\boldsymbol{A}_1\boldsymbol{u}_0 + \frac{T}{2}\left(I+\frac{\boldsymbol{A}_0}{w}\right)\boldsymbol{A}_2\boldsymbol{u}_0 - \frac{1-e^{-Tw}}{2c'w^2}\boldsymbol{A}_0\boldsymbol{A}_2\boldsymbol{u}_0.$$

*Proof.* We have $\mathbf{L}_2 := \mathbf{L}_2^1 + \mathbf{L}_2^2$, with

$$\mathbf{L}_2^1 = \int_0^T e^{(T-s)\boldsymbol{A}_0} f(s)\boldsymbol{A}_1\int_0^s e^{(s-\tau)\boldsymbol{A}_0} f(\tau)\boldsymbol{A}_1 e^{\tau\boldsymbol{A}_0} d\tau ds \tag{5.36}$$

$$\mathbf{L}_2^2 = \int_0^T e^{(T-s)\boldsymbol{A}_0} f(s)\boldsymbol{A}_2 e^{s\boldsymbol{A}_0} ds. \tag{5.37}$$

One has

$$\int_0^t e^{(t-s)\boldsymbol{A}_0} f(s)\boldsymbol{A}_1\int_0^s e^{(s-\tau)\boldsymbol{A}_0} f(\tau)\boldsymbol{A}_1 e^{\tau\boldsymbol{A}_0} d\tau ds$$
$$= \int_0^t f(s)\left(\left(I+\frac{\boldsymbol{A}_0}{w}\right) - e^{ws}e^{-wt}\frac{\boldsymbol{A}_0}{w}\right)\boldsymbol{A}_1\int_0^s f(\tau)\left[\left(I+\frac{\boldsymbol{A}_0}{w}\right)\boldsymbol{A}_1\left(I+\frac{\boldsymbol{A}_0}{w}\right)\right.$$
$$\left. -e^{w\tau}e^{-ws}\frac{\boldsymbol{A}_0}{w}\boldsymbol{A}_1\left(I+\frac{\boldsymbol{A}_0}{w}\right) - e^{-w\tau}\left(I+\frac{\boldsymbol{A}_0}{w}\right)\boldsymbol{A}_1\frac{\boldsymbol{A}_0}{w} + e^{-ws}\frac{\boldsymbol{A}_0\boldsymbol{A}_1\boldsymbol{A}_0}{w^2}\right]d\tau ds$$
$$= \int_0^t f(s)\int_0^s f(\tau)d\tau ds\left(I+\frac{\boldsymbol{A}_0}{w}\right)\boldsymbol{A}_1\left(I+\frac{\boldsymbol{A}_0}{w}\right)\boldsymbol{A}_1\left(I+\frac{\boldsymbol{A}_0}{w}\right)$$
$$-\int_0^t f(s)e^{ws}\int_0^s f(\tau)d\tau ds\frac{e^{-wt}\boldsymbol{A}_0\boldsymbol{A}_1}{w}\left(I+\frac{\boldsymbol{A}_0}{w}\right)\boldsymbol{A}_1\left(I+\frac{\boldsymbol{A}_0}{w}\right)$$

$$
\begin{aligned}
&-\int_0^t f(s)\int_0^s f(\tau)e^{-w(s-\tau)}d\tau ds\left(I+\frac{\boldsymbol{A}_0}{w}\right)\boldsymbol{A}_1\frac{\boldsymbol{A}_0}{w}\boldsymbol{A}_1\left(I+\frac{\boldsymbol{A}_0}{w}\right)\\
&+\int_0^t f(s)\int_0^s f(\tau)e^{w\tau}d\tau ds e^{-wt}\frac{\boldsymbol{A}_0}{w}\boldsymbol{A}_1\frac{\boldsymbol{A}_0}{w}\boldsymbol{A}_1\left(I+\frac{\boldsymbol{A}_0}{w}\right)\\
&-\int_0^t f(s)\int_0^s f(\tau)e^{-w\tau}d\tau ds\left(I+\frac{\boldsymbol{A}_0}{w}\right)\boldsymbol{A}_1\left(I+\frac{\boldsymbol{A}_0}{w}\right)\boldsymbol{A}_1\frac{\boldsymbol{A}_0}{w}\\
&+\int_0^t f(s)\int_0^s f(\tau)e^{-w(\tau-s)}d\tau ds e^{-wt}\frac{\boldsymbol{A}_0}{w}\boldsymbol{A}_1\left(I+\frac{\boldsymbol{A}_0}{w}\right)\boldsymbol{A}_1\frac{\boldsymbol{A}_0}{w}\\
&+\int_0^t f(s)e^{-ws}\int_0^s f(\tau)d\tau ds\left(I+\frac{\boldsymbol{A}_0}{w}\right)\boldsymbol{A}_1\frac{\boldsymbol{A}_0}{w}\boldsymbol{A}_1\frac{\boldsymbol{A}_0}{w}\\
&-\int_0^t f(s)\int_0^s f(\tau)d\tau ds e^{-wt}\frac{\boldsymbol{A}_0}{w}\boldsymbol{A}_1\frac{\boldsymbol{A}_0}{w^2}\boldsymbol{A}_1\boldsymbol{A}_0.
\end{aligned}
$$

As a result, we can compute $\mathbf{L}_2$ and use equalities (5.14)-(5.24):

$$
\begin{aligned}
\mathbf{L}_2^1 =& \int_0^T f(s)\left(\left(I+\frac{\boldsymbol{A}_0}{w}\right)-e^{ws}e^{-wT}\frac{\boldsymbol{A}_0}{w}\right)\boldsymbol{A}_1\int_0^s f(\tau)\left[\left(I+\frac{\boldsymbol{A}_0}{w}\right)\boldsymbol{A}_1\left(I+\frac{\boldsymbol{A}_0}{w}\right)\right.\\
&\left.-e^{w\tau}e^{-ws}\frac{\boldsymbol{A}_0}{w}\boldsymbol{A}_1\left(I+\frac{\boldsymbol{A}_0}{w}\right)-e^{-w\tau}\left(I+\frac{\boldsymbol{A}_0}{w}\right)\boldsymbol{A}_1\frac{\boldsymbol{A}_0}{w}+e^{-ws}\frac{\boldsymbol{A}_0\boldsymbol{A}_1\boldsymbol{A}_0}{w^2}\right]d\tau\\
=& -\int_0^T f(s)e^{ws}\int_0^s f(\tau)d\tau ds\frac{e^{-wT}\boldsymbol{A}_0\boldsymbol{A}_1}{w}\left(I+\frac{\boldsymbol{A}_0}{w}\right)\boldsymbol{A}_1\left(I+\frac{\boldsymbol{A}_0}{w}\right)\\
&-\int_0^T f(s)\int_0^s f(\tau)e^{-w(s-\tau)}d\tau ds\left(I+\frac{\boldsymbol{A}_0}{w}\right)\boldsymbol{A}_1\frac{\boldsymbol{A}_0}{w}\boldsymbol{A}_1\left(I+\frac{\boldsymbol{A}_0}{w}\right)\\
&+\int_0^T f(s)\int_0^s f(\tau)e^{w\tau}d\tau ds e^{-wT}\frac{\boldsymbol{A}_0}{w}\boldsymbol{A}_1\frac{\boldsymbol{A}_0}{w}\boldsymbol{A}_1\left(I+\frac{\boldsymbol{A}_0}{w}\right)\\
&-\int_0^T f(s)\int_0^s f(\tau)e^{-w\tau}d\tau ds\left(I+\frac{\boldsymbol{A}_0}{w}\right)\boldsymbol{A}_1\left(I+\frac{\boldsymbol{A}_0}{w}\right)\boldsymbol{A}_1\frac{\boldsymbol{A}_0}{w}\\
&+\int_0^T f(s)\int_0^s f(\tau)e^{-w(\tau-s)}d\tau ds e^{-wT}\frac{\boldsymbol{A}_0}{w}\boldsymbol{A}_1\left(I+\frac{\boldsymbol{A}_0}{w}\right)\boldsymbol{A}_1\frac{\boldsymbol{A}_0}{w}\\
&+\int_0^T f(s)e^{-ws}\int_0^s f(\tau)d\tau ds\left(I+\frac{\boldsymbol{A}_0}{w}\right)\boldsymbol{A}_1\frac{\boldsymbol{A}_0}{w}\boldsymbol{A}_1\frac{\boldsymbol{A}_0}{w}\\
=& \frac{3T}{8\pi c'}N(T,-w)\frac{\boldsymbol{A}_0\boldsymbol{A}_1}{w}\left(I+\frac{\boldsymbol{A}_0}{w}\right)\boldsymbol{A}_1\left(I+\frac{\boldsymbol{A}_0}{w}\right)\\
&-\frac{1}{c}\left(w\left(\frac{T}{2\pi}\right)^2\frac{T}{2}+\frac{T}{2\pi}N(T,-w)\right)\left(I+\frac{\boldsymbol{A}_0}{w}\right)\boldsymbol{A}_1\frac{\boldsymbol{A}_0}{w}\boldsymbol{A}_1\left(I+\frac{\boldsymbol{A}_0}{w}\right)\\
&\frac{3T}{8\pi c'}N(T,-w)\frac{\boldsymbol{A}_0}{w}\boldsymbol{A}_1\frac{\boldsymbol{A}_0}{w}\boldsymbol{A}_1\left(I+\frac{\boldsymbol{A}_0}{w}\right)\\
&+\frac{3T}{8\pi c'}N(T,-w)\left(I+\frac{\boldsymbol{A}_0}{w}\right)\boldsymbol{A}_1\left(I+\frac{\boldsymbol{A}_0}{w}\right)\boldsymbol{A}_1\frac{\boldsymbol{A}_0}{w}\\
&+\frac{1}{c}\left(-w\left(\frac{T}{2\pi}\right)^2\frac{T}{2}+\frac{T}{2\pi}N(T,w)\right)e^{-wT}\frac{\boldsymbol{A}_0}{w}\boldsymbol{A}_1\left(I+\frac{\boldsymbol{A}_0}{w}\right)\boldsymbol{A}_1\frac{\boldsymbol{A}_0}{w}\\
&+\frac{3T}{8\pi c'}N(T,-w)\left(I+\frac{\boldsymbol{A}_0}{w}\right)\boldsymbol{A}_1\frac{\boldsymbol{A}_0}{w}\boldsymbol{A}_1\frac{\boldsymbol{A}_0}{w}.
\end{aligned}
$$

Therefore

$$
\begin{aligned}
\mathbf{L}_2^1\mathbf{u}_0 =& \frac{3T}{8\pi c'}N(T,-w)\frac{\boldsymbol{A}_0\boldsymbol{A}_1}{w}\left(I+\frac{\boldsymbol{A}_0}{w}\right)\boldsymbol{A}_1\mathbf{u}_0-\frac{1}{c}\left(w\left(\frac{T}{2\pi}\right)^2\frac{T}{2}+\frac{T}{2\pi}N(T,-w)\right)\left(I+\frac{\boldsymbol{A}_0}{w}\right)\boldsymbol{A}_1\frac{\boldsymbol{A}_0}{w}\boldsymbol{A}_1\mathbf{u}_0\\
&+\frac{3T}{8\pi c'}N(T,-w)\frac{\boldsymbol{A}_0}{w}\boldsymbol{A}_1\frac{\boldsymbol{A}_0}{w}\boldsymbol{A}_1\mathbf{u}_0.
\end{aligned}
$$

and

$$\begin{aligned}
\mathbf{L}_2^2\mathbf{u}_0 &= \int_0^T e^{(T-s)\boldsymbol{A}_0}\boldsymbol{A}_2 f^2(s) e^{s\boldsymbol{A}_0} ds\mathbf{u}_0 \\
&= \frac{1}{2}\int_0^T \left(I + \frac{\boldsymbol{A}_0}{w} - e^{-Tw}e^{sw}\frac{\boldsymbol{A}_0}{w}\right)\left(1-\cos\frac{4\pi s}{T}\right) ds\boldsymbol{A}_2\mathbf{u}_0 \\
&= \frac{1}{2}\left[T\left(I+\frac{\boldsymbol{A}_0}{w}\right) - \frac{1-e^{-Tw}}{c'w^2}\boldsymbol{A}_0\right]\boldsymbol{A}_2\mathbf{u}_0 \\
&= \frac{T}{2}\left(I+\frac{\boldsymbol{A}_0}{w}\right)\boldsymbol{A}_2\mathbf{u}_0 - \frac{1-e^{-Tw}}{2c'w^2}\boldsymbol{A}_0\boldsymbol{A}_2\mathbf{u}_0.
\end{aligned}$$

□
